# Optimal control of gene regulatory networks for morphogen-driven tissue patterning

**DOI:** 10.1101/2022.07.26.501519

**Authors:** A. Pezzotta, J. Briscoe

## Abstract

The organised generation of functionally distinct cell types in developing tissues depends on establishing spatial patterns of gene expression. In many cases, this is directed by spatially graded chemical signals – known as *morphogens*. In the influential “French Flag Model”, morphogen concentration is proposed to instruct cells to acquire their specific fate. However, this mechanism has been questioned. It is unclear how it produces timely and organised cell-fate decisions, despite the presence of changing morphogen levels, molecular noise and individual variability. Moreover, feedback is present at various levels in developing tissues introducing dynamics to the process that break the link between morphogen concentration, signaling activity and position. Here we develop an alternative approach using optimal control theory to tackle the problem of morphogen-driven patterning. In this framework, intracellular signalling is derived as the control strategy that guides cells to the correct fate while minimizing a combination of signalling levels and the time taken. Applying this approach demonstrates its utility and recovers key properties of the patterning strategies that are found in experimental data. Together, the analysis offers insight into the design principles that produce timely, precise and reproducible morphogen patterning and it provides an alternative framework to the French Flag paradigm for investigating and explaining the control of tissue patterning.

## INTRODUCTION

Embryogenesis depends on positioning functionally distinct types of cells in the right place and proportions, at the right time in a developing tissue. In many cases, the arrangement of differentiating cells is guided by chemical signals (usually termed *morphogens*). Emanating from a localised source, a morphogen spreads across a field of cells to form a gradient, hence cells at different positions are exposed to different levels of the morphogen [1]. In the influential “French Flag Model” cells are proposed to read the gradient, such that the local signal concentration instructs position-dependent cell fate [2]. It has become apparent, however, that morphogen concentration alone is insufficient to explain the interpretation of morphogen gradients. In many tissues, morphogen gradients are dynamic and there is no simple relationship between morphogen concentration and position within the tissue [3, 4]. It is also unclear how a simple gradient mechanism would allow timely and accurate cell-fate decisions, despite the presence of molecular noise and individual variability.

The interpretation of the morphogen signal involves gene regulatory networks (GRNs) in responding cells [4]. These comprise the intracellular signalling pathways of the morphogens and the downstream transcriptional responses and are central to transforming the continuous spatio-temporal input of morphogen signalling into discrete cell fates. Regulatory interactions between components of these networks appear to perform the equivalent of an analogue-to-digital conversion [4–7]. GRNs have also been proposed to contribute to the accuracy and reproduciblity of patterning in presence of intracellular noise [8–10]. Moreover, non-linearities and feedback within the GRN can confer multi-stability, memory and hysteresis to cellular decision-making. A consequence of this is that cell fate depends not only on the *levels* of signals and effectors, but also on their *temporal* features. Taken together, the complexity of interactions within the GRN can produce rich dynamics in the signalling and gene expression in developing tissues. Understanding the origin and function of these dynamics offers insight into patterning. Moreover, the interplay between morphogen gradient and GRN allow cells to actively contribute to morphogen signalling, rather than being simply “instructed” by the gradient. This highlights the need for alternative paradigms to the French Flag model, in which the GRN plays a complementary and equally important role to the morphogen, to frame questions about morphogen activity.

The dorso-ventral patterning of the developing vertebrate neural tube is a well-established example of a morphogen-patterned tissue [4, 11]. In the ventral neural tube, the secreted morphogen Sonic Hedgehog (Shh), produced from the notochord and floor plate, which are located at the ventral pole, forms a ventral to dorsal gradient [12]. Binding of Shh to its receptor Patched1 (Ptch1) releases the inhibition of downstream signalling and leads to the conversion of the transcriptional effectors – the Gli family of proteins – from their repressor to their activator forms. The Gli proteins regulate the expression of a set of transcription factors, which include members of the Nkx, Olig, Pax and Irx families. This comprises the neural tube GRN. Interactions between intracellular signalling and the transcriptional network, generates a dynamic response of Gli activity to varying amounts of Shh and produces a sequence of genetic toggle switches that generate distinct gene expression states over time [3, 13]. Feedback leads to the desensitisation of cells to the morphogen signal [12, 14–16], resulting in adaption in Gli activity [16]. Similar effects of negative feedback have been observed for many signalling pathways, but its function and implications for morphogen-dependent pattern formation remains unclear.

Dynamical systems theory provides a framework to describe the activity of morphogens and GRNs. The behaviour produced by such models can often be represented geometrically as a dynamical landscape. This provides an intuitive description of cell-fate decisions that corresponds to the idea of an “epigenetic landscape” proposed by Waddington [17]. In this view, the developmental trajectory of a cell is analogous to a particle rolling on an undulating landscape, where valleys and watersheds represent fates and decision points, respectively. Morphogens can be thought of as tilting the landscape in such a way that the valleys can be made deeper, shallower or disappear altogether. In this way the morphogen controls the terrain and hence the valley a cell enters. Although originally introduced as a pictorial representation of development, this idea has been used to develop quantitative methods that reproduce key features of gene regulatory networks and make predictions about the effect of signals [18–20]. Nevertheless, it remains a challenge to construct landscape models that incorporate knowledge of signals and GRNs. How is the landscape modified by an external signal and how feedback mechanisms be incorporated? How can experimentally inferred landscapes give insights into the signalling dynamics?

Here, we set out to develop a framework to understand the intracellular signalling strategies used by cells to interpret a morphogen signal. Are there design principles to the signalling pathways that contribute to timely, precise and accurate morphogen controlled tissue patterning? What role does feedback play and does this result in a trade-off between speed, accuracy and robustness of the pattern formation? To this end, we cast the morphogen- driven patterning process as an optimal control problem, where a trade-off is sought to minimise the distance from target and the control employed. The optimization allows the activity of signalling effectors to be a function of both extracellular signal and target genes within the GRN. This function, can be considered a model of the signalling pathway which accounts for the feedback loops within it and from the GRN.

We first applied this approach to a Waddingtonlandscape model representing a genetic toggle switch – where analytical treatment is possible. We then extended the analysis to a dynamical-system model describing gene regulation in ventral neural tube progenitors. We show that desensitisation of the signalling pathway to morphogen emerges as a means to minimize control inputs in the context of multi-stability. The approach discovers morphogen patterning strategies that are widely used in biological systems and suggests an explanation for these strategies. Using this optimal control framework places morphogens and GRNs on the same footing, each playing complementary roles as parts of a whole decision-making unit. In this sense, the approach provides an alternative framework to the French Flag paradigm.

## RESULTS

### Dynamical systems and optimal control approach to cell-fate decisions

The dynamics of gene regulation and cell-fate decisions can be described using a Langevin equation

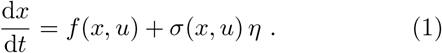

where *x* is the set of concentrations of the components of network, *u* is a set of inputs or control variables. The functions *f* and *σ* are the drift and the strength of the noise, respectively (*η* is a standard white noise). In general, the noise term has a multiplicative form, which accounts for stochasticity that arises not only from external disturbances but also from the finite number copy number of each species in the network [21]. The drift and noise functions *f* and *σ* can incorporate mechanistic knowledge of the regulatory logic of the network and the effect of morphogen signalling, for instance, transcriptional control via binding/unbinding of transcription factors to their respective regulatory elements and cooperative and competitive effects [13, 22, 23].

The dynamical systems that result from representing GRNs in this way are generally non-linear and may operate in multi-stable regimes. The input *u* can substantially change the dynamics of the network, altering the position of attractors (stable states) and saddle nodes (decision points). Moreover, the attractor reached by a system depends on the full past history of the inputs. This can be seen, for example, in the neural progenitor GRN [13], where the input *u* comprises the activating and repressing forms of the morphogen regulated Gli effectors (Fig. 1 (a) and (b)). The behaviour of such systems can be visualised as a dynamical landscape with valleys representing the stable states of the network and signals tilting the landscape to determine which valleys are accessible or inaccessible. The dynamical system function *f* is thus given by the gradient of the landscape, *V*, parametrically dependent on the effector *u*. This approach has been used to reproduce the qualitative features of GRNs as well as to predict patterning processes in embryos [18, 19] and proportions of cell types in differentiation protocols [20].

**FIG. 1.**
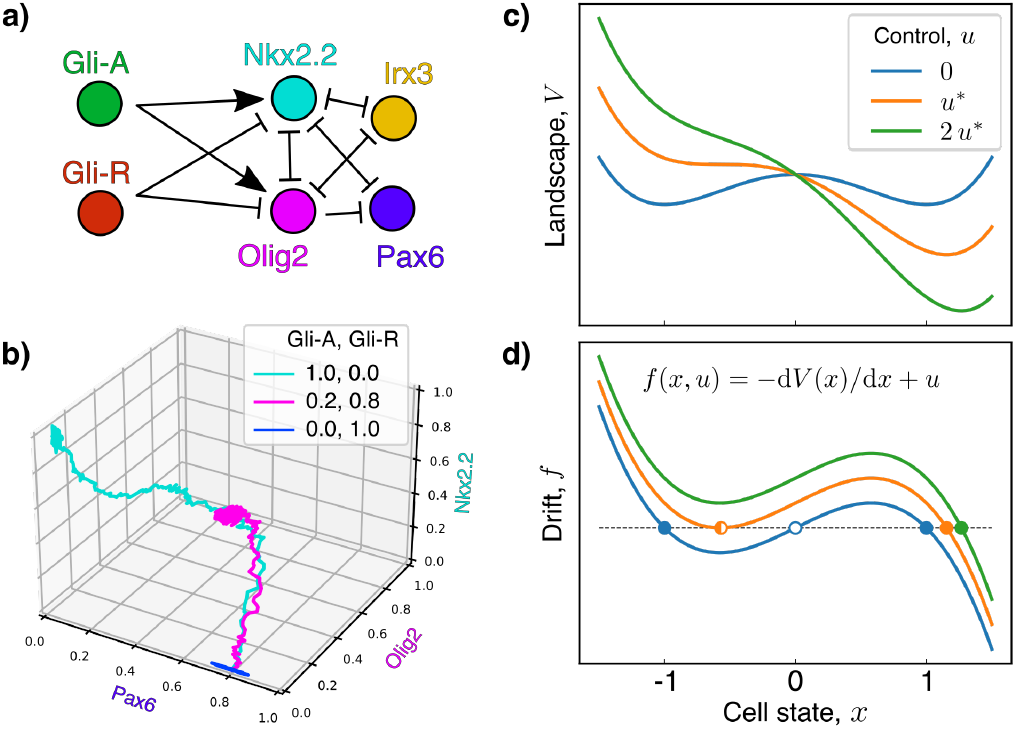
External input changes the stability properties of the dynamical system. (a) We consider a model of gene regulation which describes the patterning dynamics in the ventral neural tube with the addition of intrinsic noise [8, 13]. (b) Different levels of the inputs Gli-A and Gli-R (see legend) result in qualitatively different trajectories in gene expression space. Starting from the same state (low Nkx2.2 and Olig2, but high Pax6 and Irx3 – the latter suppressed in the 3D plot), the trajectories end in different stable fixed points. (c) In the Waddington-landscape picture, cell-fate decisions can be thought of as a drive towards different possible minima of a potential landscape, the “depth” of which are controlled by external signals (*u*) that “tilt” the landscape. (d) In this analogy, cell-fates are the stable fixed points of the corresponding dynamical system – the minima of the landscape (full circles). Varying external inputs changes the dynamical properties of the system, by creating and destroying attractors and fixed points; for instance, a saddle-node bifurcation corresponds to the coalescence of a stable fixed point with an unstable fixed point (empty circle).

Given this dynamical systems view of patterning, how does the signalling input to a GRN generate a sufficiently precise pattern in a developmentally relevant time period? To address this we recast patterning as an optimisation problem and ask what sort of signal input is necessary to produce precise, reliable and timely cell-fate decisions. The framework that naturally deals with these types of problems is optimal control theory. We are faced with the task of choosing a dynamic signalling regime *u* (here referred to as *control)* that minimizes the average of a cost accumulated along the trajectory plus a cost determined by the distance from the target at the termination of the decision task – in the cell-fate decision case, a differentiation event. This can be expressed in terms of a cost rate 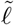 (or running cost) that gives a measure of the instantaneous performance, along with a terminal cost *Q.* We construct the function 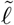 to measure how far gene expression deviates from its target (via a function *q*, which is minimum at the target) and how much control is exerted in the process, e.g. by adding a term quadratic in *u* weighted by a parameter *ϵ*; the terminal cost *Q* is also chosen to measure the distance from the target, and is here assumed to be identical to *q* up to a unit time constant. In summary, we express the cost

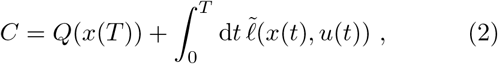

and seek a function *u* minimising its mean over realisations of the dynamics in Eq. (1). Here, *T* is the random time of differentiation, which is assumed to be exponentially distributed, with mean *τ* – or, equivalently, to occur at any time with uniform probability rate *τ*^-1^.

From the point of view of decision making, and therefore planning, the constant rate of differentiation assigns more weight to more imminent events, while discounting those further away in the future (see SI, Sec. SI-1b). As shown in SI, Sec. SI-1c, the minimisation of the cost in Eq. (2) is equivalent to that of

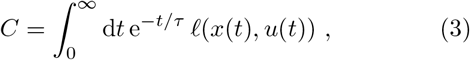

where 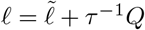. This form of the cost explicitly expresses the notion of future discounting. For these cost functions, the conditions for optimality acquire the form of differential equations, and yield the optimal *u* in the form of feedback control, *u*\*(*t*) = *φ*\*(*x*(*t*)) (see Sec. in Methods and SI). This framework is particularly relevant in the context of the control of gene expression in a cell, where aspects of the signal transduction pathway and the signal effector can be under the control of the transcription factors in the GRN (Fig. 2 (b)). When the optimality equations cannot be solved analytically or numerically, approximate solutions can be found via techniques such as reinforcement learning (RL) [24]. Solving for the optimal control *u*\*, yields optimal feedback designs and can shed light on the functional role of observed feedback mechanisms.

**FIG. 2.**
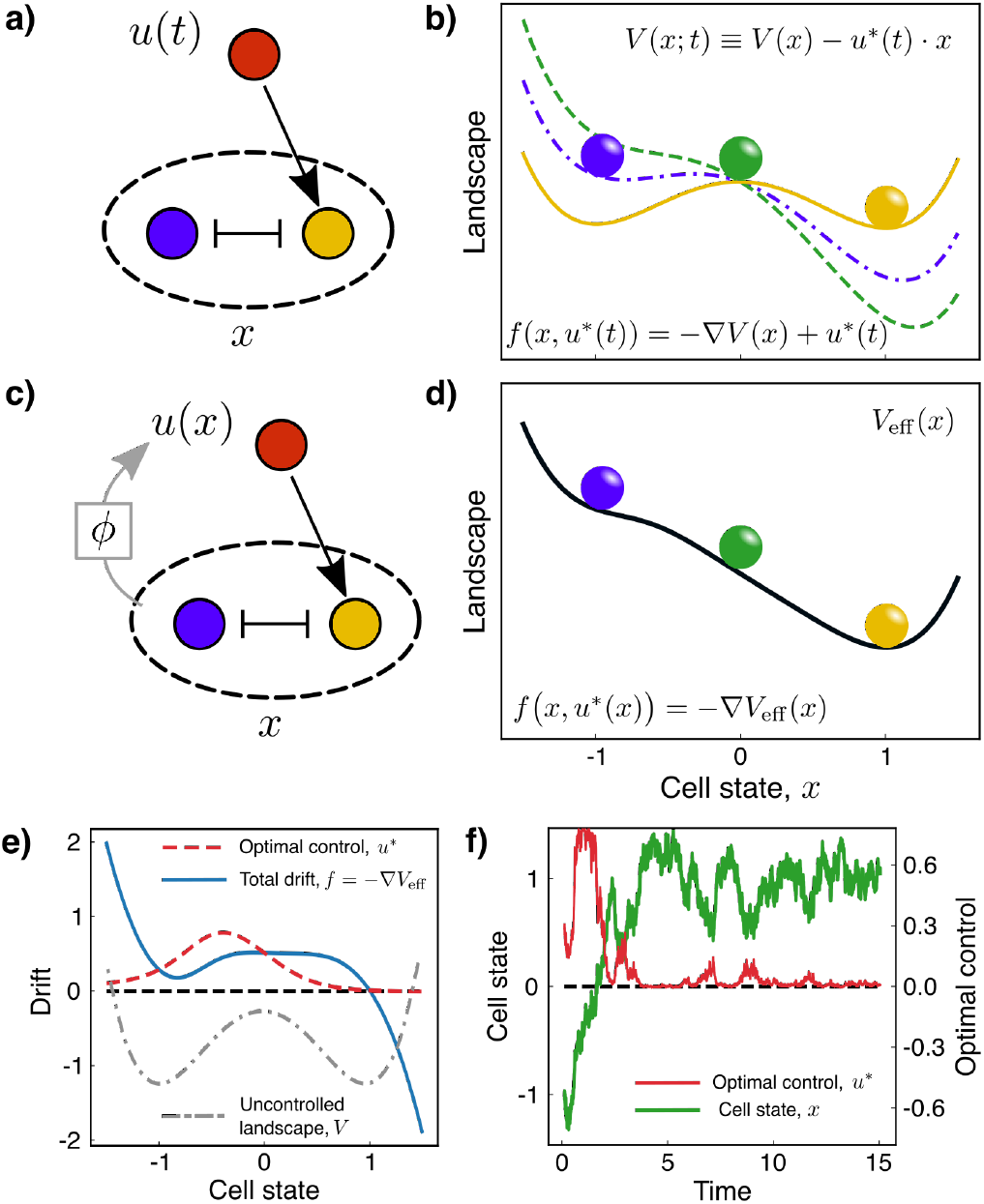
Optimal control representation of a Waddington landscape. (a) A GRN for a simple toggle-switch network with two genes can be dynamically controlled to reach a target state by explicitly defining a signalling protocol *u*(*t*) (open-loop control). (b) In the Waddington-landscape picture, we can think of the external control as “tilting” the landscape over time; the coloured lines represent the instantaneous landscape felt by the “particle” of the same colour. (c) Alternatively, the signal can be placed under control of the target genes through a feedback function *ϕ.* This results in closed-loop, or feedback, control. (d) The optimal closed-loop control is incorporated into a “static” effective landscape, describing the dynamical properties of the signalling and GRN system as a *whole.* (e) The solution for the optimal control (dashed red line) exhibits adaptation near the target, when this corresponds to a stable fixed point of the uncontrolled landscape (dashed-dotted grey line, not in scale). (f) This can also be seen in a sample trajectory of the dynamics of a cell (green line), where the control (red line) is switched off after an initial transient, and is activated only to prevent large fluctuations away from the target. For (e) and (f), the parameters used are *D* = 0.10, *τ* =10 and *ϵ* = 10.

### Controlling the epigenetic landscape of a genetic switch

In order to illustrate this method, and to understand the parameters of the cost function, we first considered a simple model for a binary cell-fate decision. A one-dimensional double-well potential *V*(*x*) with minima at ±1, which correspond to two possible cells fates (see SI, Sec. SI-1). In this example, the noise is modelled as additive and independent of control, i.e. 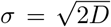, with constant *D*. We model morphogen signaling as a drift contribution *u*, which “tilts” the landscape, *V*(*x,u*) = *V*(*x*) – *u* · *x* (Fig. 2 (c)). We then seek to find the control protocol *u* (the dynamics of signal) that drives a cell from state *x* = –1 to the state *x* = 1 in the optimal way, i.e. minimizing the combination of how far the cell is from its target and the amount of control exerted to accomplish this (see SI, Eq. (S2) and (S16)).

In this model, an exact solution of the optimality equations can be found with numerical methods. The resulting optimal control protocol leads to adaptive dynamics: high levels of control are necessary to leave the initial attractor, then as the system approaches the target attractor, the amount of control is minimal, and only required to prevent noise from reversing the transition (Fig. 2 (f), and Fig. S1). From this example we see that the optimal solution minimises control by taking advantage of the multi-stability built in the system.

The linearity of the dynamical system with respect to *u* and the quadratic cost for control, means that the optimally controlled drift can be expressed as the negative gradient of a landscape function *V*_eff_. This represents a combination of the original landscape V and the optimal cost expected to be paid from a given state *x* (the cost- to-go function, see Methods). Thus, rather than thinking of the control as tilting the landscape over time, it can be incorporated into a new landscape that describes the system as a whole (Fig. 2 (d)). This observation suggests that the inverse problem might provide insight into the function of feedback mechanisms in cell-fate decisions: given experimental observations and a landscape associated with the underlying GRN, it might be possible to distinguish the contributions of the controlled system (the GRN) from the feedback mechanisms (Fig. 2 (e)).

This example also provides intuition into the effect of the differentiation rate – equivalently, of discounting cost over time. What is the optimal behaviour of the system before a cell differentiates?

At one limit, when the differentiation rate is high, *τ* ≃ 1 (in units of the overall time-scale of the system), and noise, *D*, is low, only imminent running costs and the terminal cost are taken into account in planning, and the optimally controlled dynamical system is bistable. This is because when the system is far from its target, a substantial reduction in the distance of the system from its target within a short time *τ* would have a very high cost for control. Therefore, the only part of the cost that the controller *can* minimize is the cost of control itself. This leads to low values of the control at every state, and the system remains within the bi-stable regime (Fig. 3 (a,c) and S1, bottom left). Such small values of *τ*, would mean that a cell only rarely reaches its target before differentiation.

**FIG. 3.**
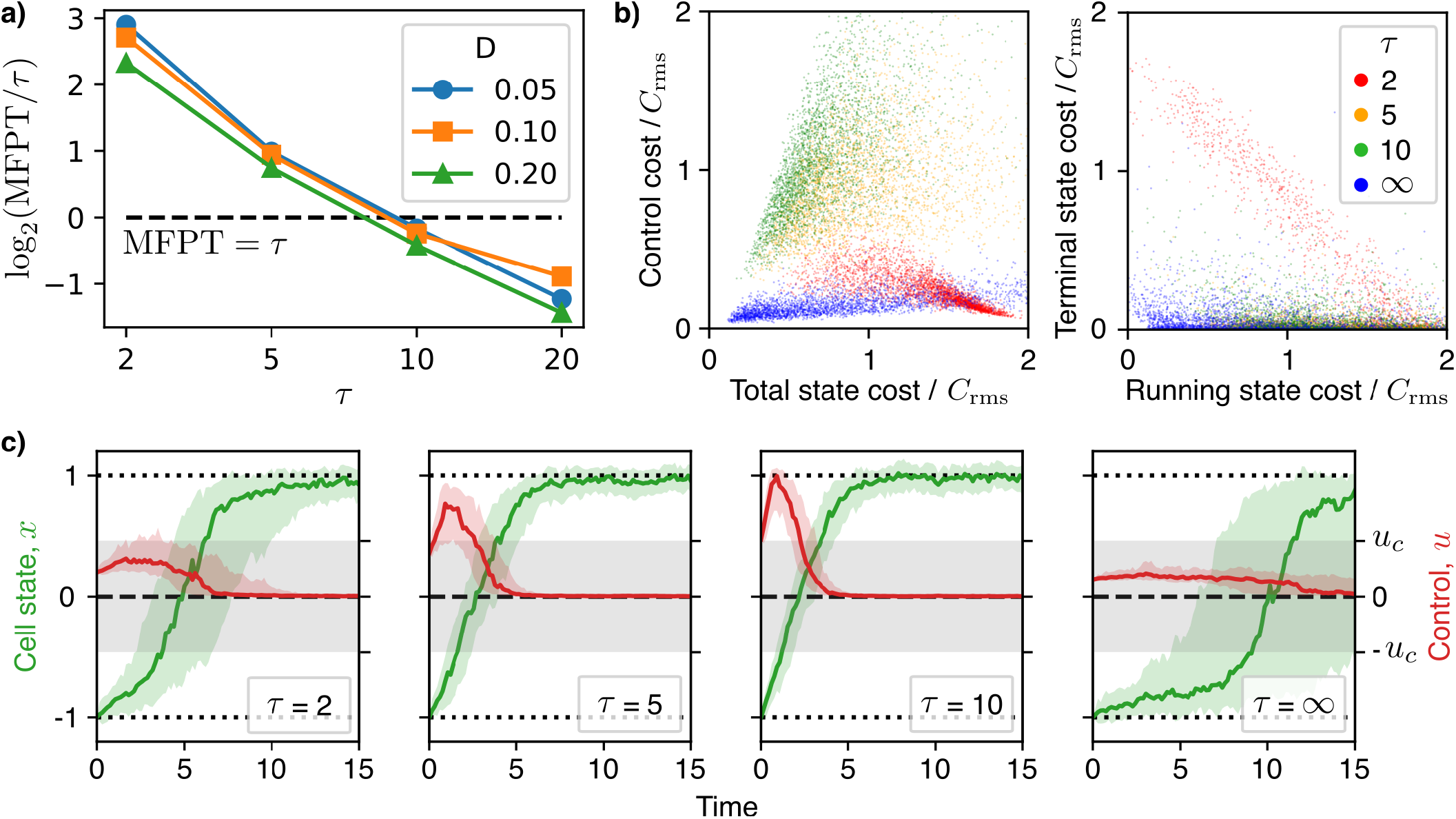
Effect of the discounting (differentiation) time *τ*. (a) The mean first passage time (MFPT) at the target *x* = 1 from *x*_0_ = −1 as a function of *τ*, from the numerical integral of the analytical formula, under the optimal control. This is shown relative to the value of *τ* on a logarithmic scale. For high (low) values of *τ*, the MFPT for the optimally controlled dynamics is far lower (higher) than *τ* itself, and decreases with the strength of the noise, *D.* (b) State and control costs from 5000 simulations for various values of τ (colour-coded). The optimal control for “small” or “large” values of *τ*, effectively minimises cost for control, while for intermediate values of *τ* a non-trivial trade-off is observed (left panel). Only for low values of *τ* ~ 1 does the terminal cost for the distance from the target have a large contribution to the overall cost (right panel). (c) Statistics of 100 samples of the dynamics for the state (green) and the control (red). Solid lines are the median values, shaded areas the 25-75 percentile. The grey shaded area highlights the values of the control variable *u* for which the controlled landscape is still bistable, i.e. between the bifurcation values ±*u_c_*. In all panels, *ϵ* = 10; in b) and c) *D* = 0.05. For intermediate values, when the MFPT is comparable to *τ*, the switch is driven by a non-trivial transient dynamics for the control, resulting from competition between control and target running costs.

Strikingly, very similar dynamics are observed in the opposite limit, when *τ* = ∞ (Fig. 3 (a,c) and S1, top left). Here, no terminal cost is paid, and the problem consists of optimising the average cost per unit time at steady state. For low *D*, when multiple stable fixed points are present (as in the case of small *u* – bistable regime), the system spends long periods of time near each of them, with rare stochastic transitions between. In SI, Sec. SI-1d, we demonstrate how the steady-state average of the cost *q* is exponentially small in *u/D*, when *D* is small: this allows very low values of *u* to yield large discrepancies between the probabilities of being in either attractor at steady state. This explains why, in such limit, it is optimal to choose *u* well within the bistability regime.

For intermediate values of *τ*, the optimally controlled dynamics are such that the time needed to perform the switch is comparable with *τ* itself. When this is the case, characteristic transient dynamics are observed: in a first phase, high levels of control are applied to the system in order to drive the transition; in a second phase, the control can be reduced to very low levels, within the bistable regime. This suggests that, in these scenarios, the optimal strategy is for the controller to apply high levels of control for a short time resulting in a lower cost from being off target for a shorter period of time (Fig. 3 (a,d)). This effect is less and less pronounced with increasing noise levels, *D*: the distribution of transition rates are controlled more and more by noise, with a smaller and smaller average transition time (Fig. 3 (a)).

By making use of a simple Waddington landscape model, this example shows how optimal control theory can make sense of adaptation as the most “parsimonious” strategy to drive a cell to a desired target, while exploiting the multi-stability of a downstream network and its stochastic dynamics. The analytical results suggest an explanation for optimal signalling in the face of varying degrees of noise and multi-stability, and for different values of differentiation rates, which set the exponentially distributed time horizon within which cell-fate decision needs to take place.

### Control of cell-fate in ventral neural progenitors

Next, we applied this optimal control approach to a GRN model that captures the patterning dynamics in the ventral region of the developing neural tube [13]. In this model noise from fluctuations in the copy number of components of the system have been introduced using the chemical Langevin equation approximation [8, 22] (Fig. 1, and reported in SI, Sec. SI-2a). The control here is a two component vector representing the activator and repressor form of the morphogen controlled Gli effectors. These directly regulate the two most ventral markers, Nkx2.2 and Olig2 (Fig. 2 (a,b)). In this case, we find an approximate solution of the optimal control equations via reinforcement learning (RL) [24]. RL provides the means to identify optimal control strategies, without knowledge of the dynamical system function *f*, by sampling states, actions (controls) and running costs (or reward signals). Here, and in the following section, we use the TD3 algorithm [25] which is a state-of-the-art RL algorithms for continuous control problems (see SI, Alg.1 for details). Using this approach we identify optimal control strategies for the system to adopt an Olig2 state or a Nkx2.2 state.

#### Algorithm 1 Twin Delayed Deep Deterministic (TD3) policy gradient for episodic tasks.

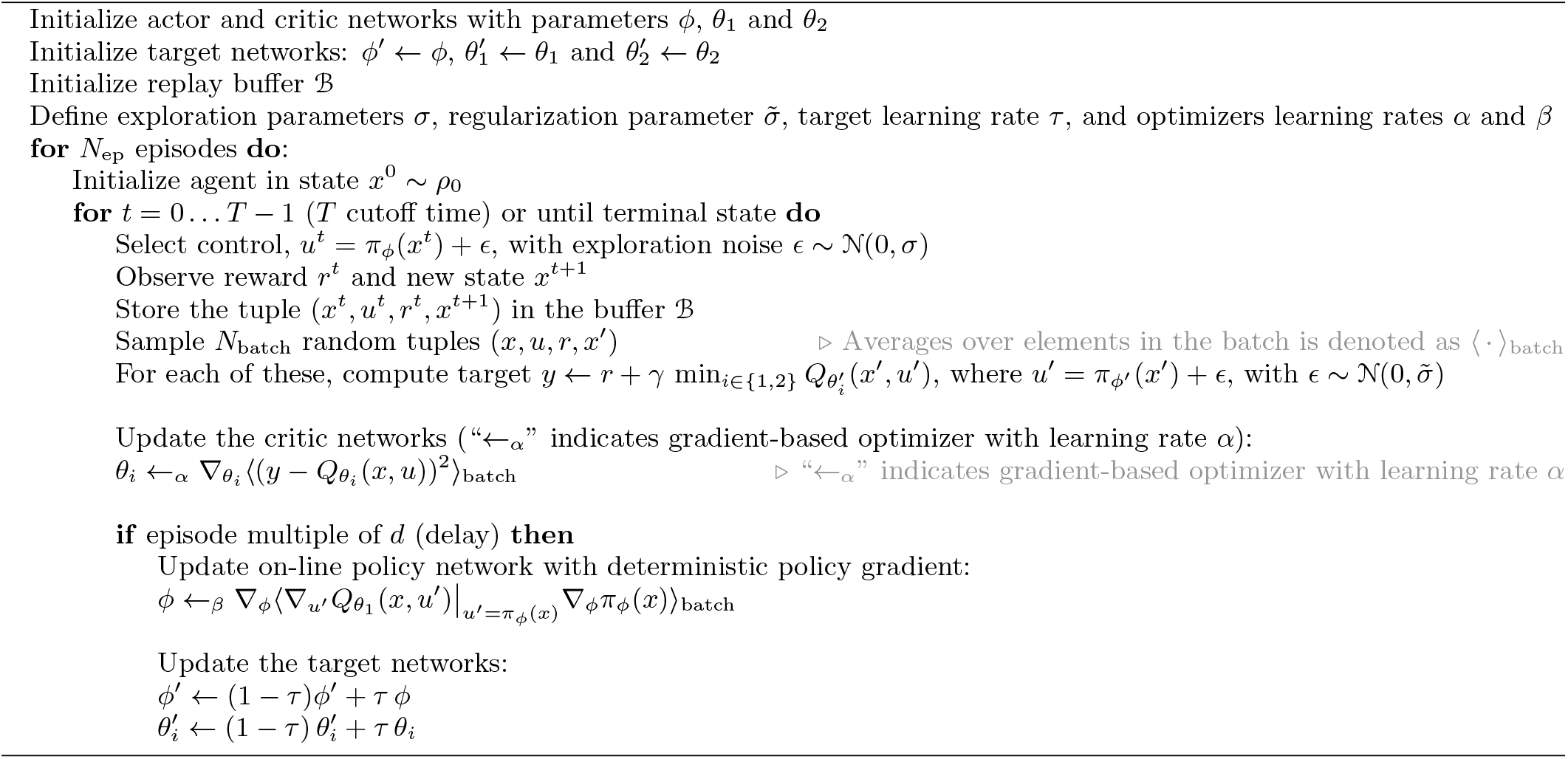

In all cases, we optimize the discounted cost function, Eq. (S16), with *τ* ≃ 5 (A.U.): this can be compared to the half-life of Nkx2.2 and Olig2, t_1/2_ ≃ 0.35 (in simulation units – see Tab. I in SI). Thus, if t_1/2_ ≃ *4h* then *τ* ≃ 2.5 days, consistent with the developmental time scales in the embryonic mouse neural tube. For both targets, the control input shows a very clear transient. Convergence of the RL algorithm to an optimal strategy in the transient is hard to achieve due to the poorer sampling of the transient configurations, resulting in run-to-run variance; however, the control strategy at steady state is consistent throughout experiments (see Fig. S3).

Acquiring and maintaining the Olig2 state requires a very high sensitivity of control with respect to Olig2 levels, which is reflected in the high variability of the repressive form of Gli effector at a population level (Fig. 4 (a)). The learnt control is such that below a threshold value of Olig2, Gli repressor is high, and above the threshold Gli repressor is low (Fig. 4 (b)). One explanation for this could be that higher levels of repressor are necessary to restrain the system from bifurcating to Nkx2.2 when levels of Olig2 are too low. This is consistent with the experimental evidence that Olig2 may provide negative feedback onto the expression of Gli3, which is the dominant repressor for Shh signaling [16, 26, 27].

**FIG. 4.**
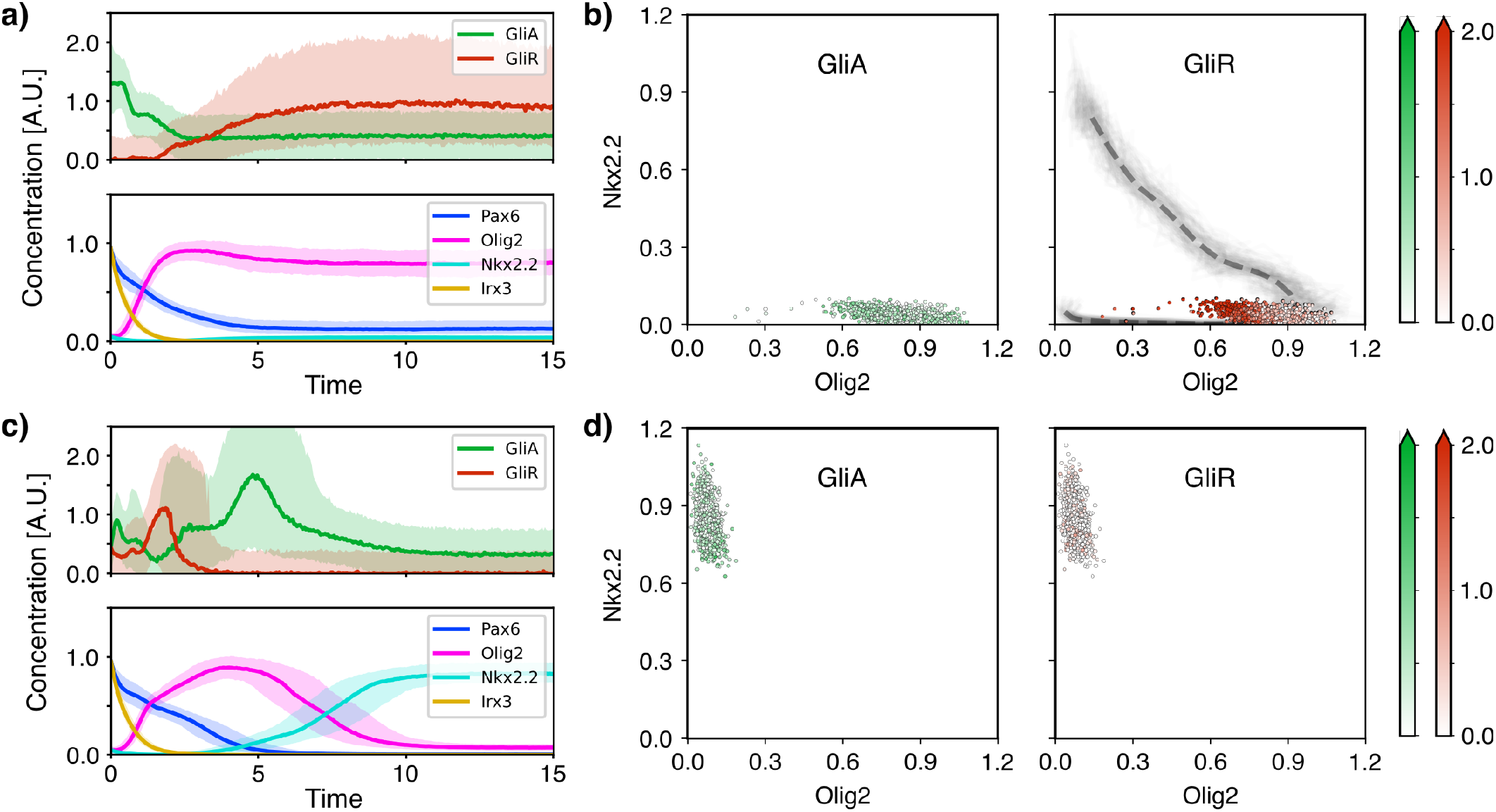
Reinforcement learning solution for the optimal control of the ventral neural tube GRN. (a) Samples of the controlled dynamics for the Olig2 target. The control *u*\*, comprising activator and repressor Gli (top panel) and the resulting gene expression dynamics (bottom panel). (b) Snapshot at steady state of the optimal control *u*\* for activator Gli (left panel) and repressor Gli (right panel) as a function of Olig2 and Nkx2.2 levels. In (c) and (d), the analogous plots, for the Nkx2.2 target. In both cases, Gli activity (relative value of activator vs repressor) is high in a first transient, and decreases over time. A negative feedback from Olig2 onto the repressor appears to be required to maintain cells in the Olig2+ state – see (b), right panel. One possibility is that this prevents the activator driving the state towards Nkx2.2+ state (the optimally controlled trajectories of panel (c) are overlaid as grey lines – the dashed grey line is the average).

This can be compared to the result for the Nkx2.2 target. Similar values for the activator form of Gli are found at steady state, but much lower values for Gli repressor are observed. The overall low levels of the effectors is also consistent with the repressive role of Nkx2.2 on Gli gene expression, as supported by experimental data [15, 16, 27]. It is notable that under the optimally controlled dynamics, a cell reaching the Nkx2.2 target must transition through the Olig2 state before acquiring Nkx2.2 expression.

### Morphogen-driven patterning

In the previous section we identified optimal control strategies independently for two target states. Here we extend the approach to identify an integrated optimal control strategies that would generate a morphogen patterned tissue comprising multiple states in response to a spatially graded morphogen signal. We then define the state of the controlled system to comprise the GRN state and the signal as subsystems.

Patterning, as an optimal control problem, can be conceived as a cooperative multi-agent task, whereby multiple cells have to reach their respective targets simultaneously, but where the shared morphogen input provides the positional information. Collectively, cells minimize a global shared cost, with the constraint that controller function – representing the signalling pathway with its feedback loops – has to be the same for all cells. The target pattern, implemented through the running cost *q*, has two boundaries that divide the tissue into three equal parts, with ventral, middle and dorsal fates corresponding to Nkx2.2, Olig2 and Pax6+/Irx3 expressing, respectively. We adapt the TD3 algorithm for the patterning task, and test it on the patterning of the ventral neural tube (see SI, Alg. 2).

#### Algorithm 2 Multi-Agent Twin Delayed Deep Deterministic (TD3) policy gradient for episodic tasks

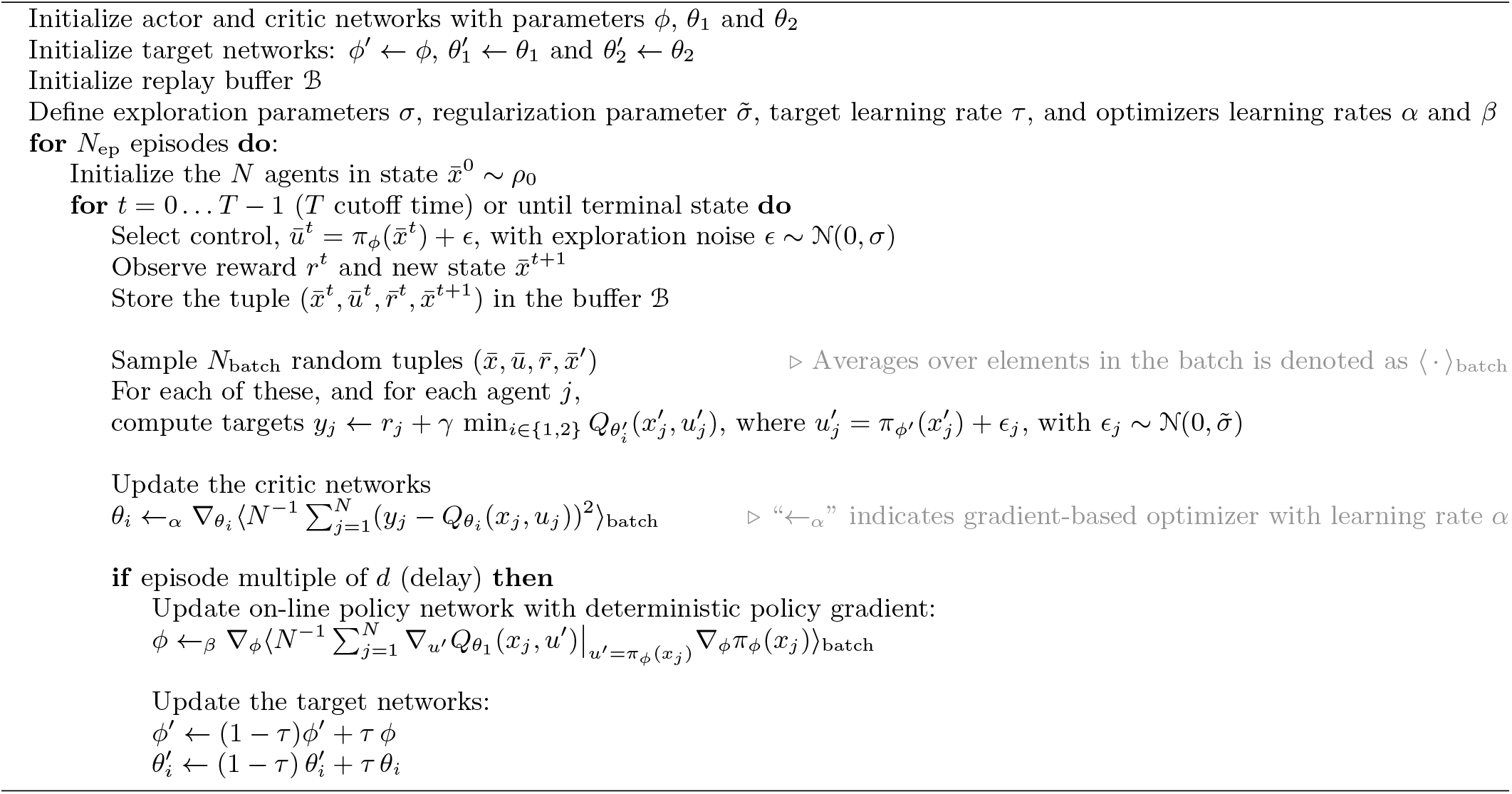

The morphogen dynamics are given by stochastic simulations of a diffusion process of independent Shh particles, while the GRN model is the same as in the previous section (details in SI, Sec. SI-2). We derive the optimality equation for this, in the ansatz of independent cells (in SI, Sec. SI-3. This ansatz can only be an approximation to the optimal solution, because the (stochastic) morphogen dynamics exhibit spatio-temporal correlations. Indeed, it works for a deterministic and static gradient – where the ansatz is exact (Fig. S4) – and can be a good approximation when the steady-state of the morphogen is reached fast compared to the GRN. A naive implementation of the independence ansatz for a “slow” morphogen fails to reproduce the target pattern, due to the increasing effect of the correlations between morphogen signals at different locations in the tissue. Nevertheless, the (ensemble) average of the morphogen signal experienced by individual cells can be expressed with independent but non-autonomous dynamics (see SI, Sec. SI-2b).

This suggested that the introduction of memory variables into the decision making may help to solve the problem, by “extracting” temporal features of the morphogen (Fig. 5 (a), and SI, Sec. SI-3 c). These variables can be thought to represent the intermediate components in the signalling cascade, such as the Shh receptor Ptch1 and the transmembrane protein Smo etc. The activity of these components in response to Shh introduce delays and persistence to the transmission of the instantaneous changes in the morphogen. The control model we introduce features more general feedback mechanisms within the signalling cascade and from the GRN species. With this extension, the algorithm is able find strategies that lead to the target pattern (Fig. 5 (b)), which we were not able to achieve without the memory variables.

**FIG. 5.**
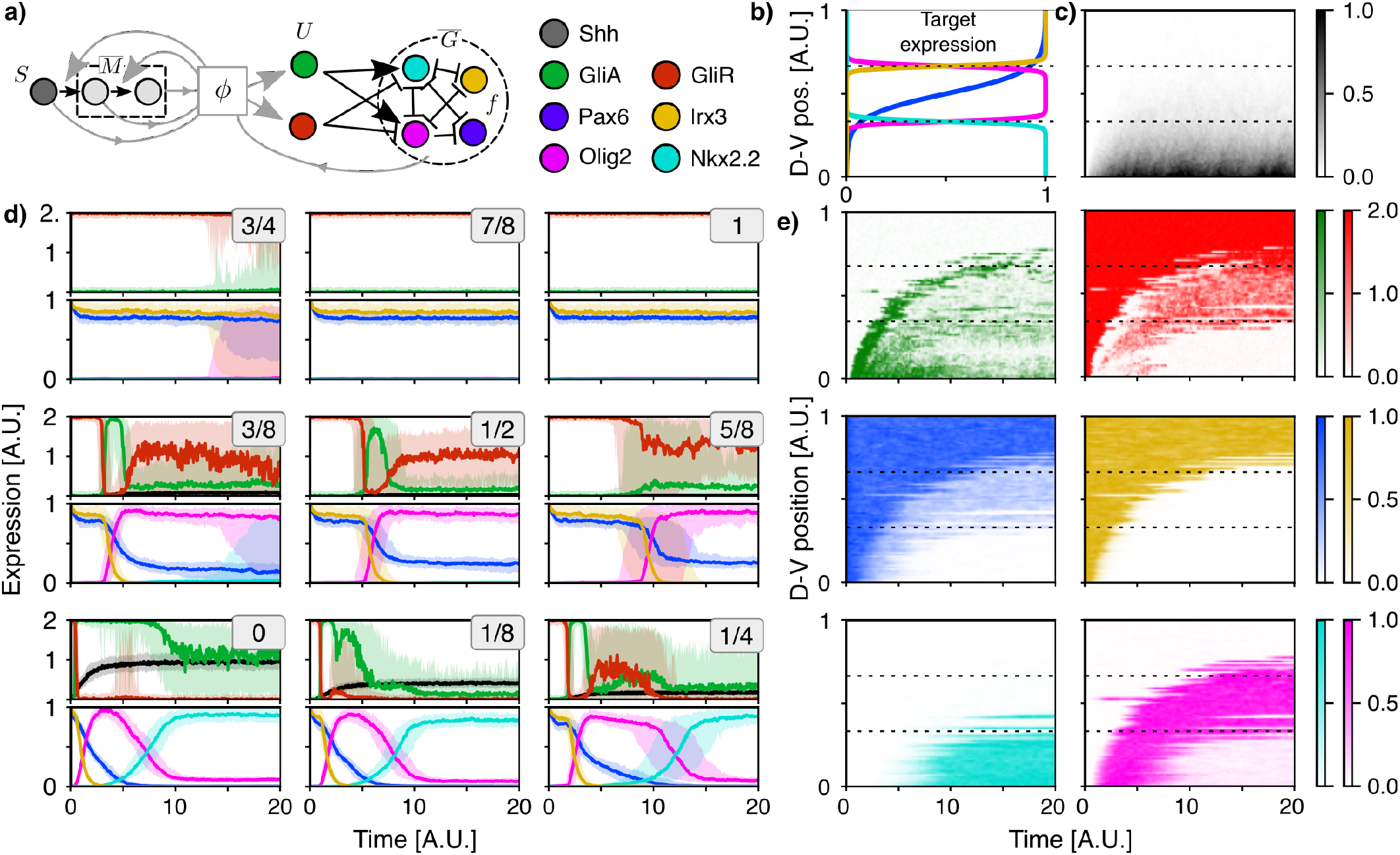
Reinforcement learning solution for the morphogen-driven patterning task. The optimal control model (a) gives the signalling effectors *U* (Gli-A/R) as a function *ϕ* of the target genes *G,* the morphogen signal *S* and memory variables *M*. The goal is to minimise a trade-off between the distance from a target gene expression profile (b) and the magnitude of the control over time. The dashed lines at 1/3 and 2/3 of the total D-V extension indicate the positions of the boundaries between target differential expression regions. The patterning process is driven by a stochastic diffusion of the Shh morphogen *S* (c). In (d), the cell-by-cell view of the dynamics averaged over 100 simulations (solid lines are the medians, and the shaded areas the 10-90 percentile, and individual panels are labelled by the D-V position of the selected cells) reveals the control strategy for each position. Similar features shown in Fig. 4 are also found here, highlighting the potential functional role of Gli repression by Olig2 and Nkx2.2 in the patterning process. In (e) a single realisation of the optimally controlled dynamics with the morphogen field as in (c).

In Fig. 5 (b), we see the average of several simulations of the tissue patterning process: at the beginning of the morphogen spread, all cells are in the initial pre-pattern (dorsal) condition. As morphogen spreads into the tissue, Olig2 and Nkx2.2 are sequentially induced ventrally, resulting in a kinematic wave of gene expression spreading from ventral to dorsal until the target pattern is reached. The pattern is then maintained. The dynamics of the effectors in individual cells (Fig. 5 (c)) share some features with those found for the single cell control (Fig. 4 (a,c)). Because the initial conditions are the same for all cells in the tissue (Pax6+/Irx3+, vanishing morphogen signal and memory variables – see SI, Sec. SI-3c), the signal levels are also the same, corresponding to the values needed to maintain cells in the dorsal state, i.e. high levels of repressor together with low levels of activator (Fig. 5 (c), top). For cells that are assigned to an Olig2+ fate, after an initial delay set by the spread of the Shh morphogen, the dynamics are remarkably similar to those found for the Olig2 target in a single cell: levels of repressor negatively correlated with Olig2 concentration and low levels of activator at steady state (Fig. 5 (c), middle). In cells acquiring an Nkx2.2+ fate we also observe a negative correlation of Gli repressor levels with Nkx2.2 (Fig. 5 (c), bottom). Thus, the learnt control strategy recovers the repressive feedback from both Olig2 and Nkx2.2 on Gli, which results in adaptive dynamics of the signalling effectors. Both of these features are supported by experimental data [15, 16, 26, 27].

## DISCUSSION

Here we used optimal control theory to develop a framework to analyse morphogen signaling strategies and identify mechanisms that produce rapid, precise and reproducible cell-fate decisions during tissue patterning in embryo development. We demonstrate that this framework can be combined with dynamical – Waddington – landscape models of cell-fate decisions to provide an optimal control representation in the form of a new landscape. Reinforcement Learning can be used to solve optimal control problems associated with signalling and cell-fate decisions and we formulate the patterning problem as a multi-agent cooperative optimal control task, in which the objective function is a measure of performance of all the cells in the tissue. By using these approaches to analyse the morphogen patterning of neural progenitors we highlight how the optimal mechanisms obtained are consistent with experimental data.

The analysis revealed that for both individual cell fate decisions and for morphogen-driven tissue patterning, adaptive signalling dynamics, which are observed experimentally *in vivo* [28], emerge as an optimal strategy in the presence of multi-stability. This suggests that signalling pathways have evolved to take advantage of the dynamical landscape that arises from the gene regulatory network. By contrast, in the celebrated French Flag model of morphogen patterning, cell fates are proposed to be instructed by morphogen concentration with the concentration viewed as being read out directly by cells [2]. While the French Flag model has been crucial for highlighting the role of morphogens in pattern formation, it does not explain the complex cellular signalling dynamics that are often observed experimentally. Moreover, it subordinates the role of the GRN to that of the extracellular signals. The optimal control perspective provides an alternative paradigm that accommodates the dynamics in signal interpretation and establishes a relationship between the control signal and the system. This provides a framework that complements dynamical systems approaches to gene regulation – where signals are externally imposed – by making signalling an integrated part of a whole decision-making unit: the cell.

The objective function includes a notion of “timing” through exponential discounting. This can be regarded as representing the tempo of development and the rate of differentiation in a tissue, which limits the amount of time that is available to the cell to integrate the signal and make a decision. We set this time to be comparable with differentiation rates and the degradation rates of the key transcription factors in the GRN [29].

Importantly, when a Waddington landscape offers a good phenomenological model of cell-fate decision, the optimal control framework provides analytical tools to “isolate” the contribution of morphogen signalling to the GRN dynamics. Practically, this could be achieved via the comparison of experimentally measured landscapes under different genetic or pharmacologic manipulations of signalling pathways [20].

There are limitations to our approach that will need to be addressed in future work. In the current formulation, the control input to the system is selected in a “reactive” way, as a function of the target genes. This rules out possible hysteresis effects in feedback mechanisms. This is partially addressed via the addition of memory variables in the morphogen-driven tissue patterning example. Yet, the signalling effectors – as a function of components of the GRN – still retain a memory-less component. This could be tackled by introducing production-degradation dynamics, where the control defines the production rates, rather than the levels. This would have the benefit of allowing the inclusion of known kinetic properties of the effectors, such as degradation rates [29]. Also, the degradation rate has been assumed independent of the cell state. The control problem solved here can be extended to cases where the terminal-time statistics depends on the state and control variables, and include optimal stopping time problems (see e.g. [30]).

From the RL perspective, the introduction of memory variables is analogous to the use of recurrent networks for modelling systems with memory [31], e.g. in partially observable environments [32, 33]. Examining this problem in the broader context of decision making in non-Markovian or non-stationary environments [34] could highlight general design principles that optimally deal with memory. It is interesting to note that the morphogen-driven patterning task can be formally regarded as a classification of signal time series: hidden in the optimally-controlled dynamics are the features of the temporal profile of the signal which can be utilised by the cell in order to make decisions. Hence the optimal control perspective provides a link between the complex computational problem of morphogen interpretation and the biological hardware available for its solution.

We did not address all possible feedback mechanisms that could be exploited by the system. For example, Shh signaling controls the expression of Shh binding proteins, such as Ptch1, Scube2 and Hhip1, that alter transport of the morphogen through the tissue [12, 14, 35]. Feedback on morphogen spread could be incorporated into the model. Indeed, the framework could be used to investigate virtually any aspect of the system. This could include, for example, control of diffusivity of signals, degradation rates of system components, or the accessibility of cis-regulatory elements and the effect of chromatin remodelling. All of which have been implicated in the interpretation of morphogen signalling [1, 8, 14].

The patterning example dealt with in this study is one in which positional information is provided by a signal external to the tissue. In other cases, symmetry is broken and patterning controlled by internally generated signals, such as in the case of organoids patterned by Turing-like mechanisms [36] Patterning, in these contexts, poses a problem of coordination by means of signalling that can be cast into a multi-agent decision making task. This, in turn, can be tackled numerically with multi-agent RL (MARL) algorithms [37, 38] or analytically via, e.g. mean-field approximation in the limit of large numbers of cells [39, 40]. Therefore, optimal control provides a framework in which to analyse these systems to investigate functional explanations for the observed signalling strategies, proportions of cell types and self-organisation of patterning.

The optimal control approach, with its focus on linking mechanisms with control, is ideally suited for the analysis of *in vitro* and synthetic systems. This could be used to design and refine signalling regimes for the directed differentiation of stem cells *in vitro* and the production of specific sets of cell types in defined proportions. An understanding of the control principles operating in biological systems will provide insight and inspiration for the construction of artificial systems and will support the use of stem cells in disease modelling and regenerative medicine.

## ACKNOWLEDGEMENTS

We are grateful to Rubèn Perez-Carrasco and Zena Hadjivasiliou and members of the lab for their constructive comments. A.P. thanks Antonio Celani for insightful discussions. A.P. was funded by the EMBO Long Term Fellowship ALTF 860-2019. This work was supported by the Francis Crick Institute, which receives its core funding from Cancer Research UK, the UK Medical Research Council and Wellcome Trust (all under FC001051). Work in the Briscoe lab is funded by the European Research Council under European Union (EU) Horizon 2020 research and innovation program grant 742138.

## METHODS

### Optimal stochastic control and its solution

Given a system with state variables *x* satisfying the controlled stochastic dynamics

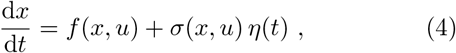

where *f* is a deterministic drift, *σ* – multiplying the standard Gaussian white noise *η* – is the magnitude of the noise and *u* represent a set of control variables, we ask what is the optimal choice of the control variables *u* over time in that minimizes the mean of a cost function

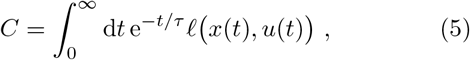

where *ℓ* is a cost per unit time (also termed running cost) associated withto the instantaneous state and control at a given time, and *τ* sets the time-scale for the exponential discount factor – defining the “far-sightedness” of the decision-maker in the estimation of the cost that is expected to be paid in the future. As we show in SI, Sec. SI-1c, optimal-control problems with terminal-state cost and uncertain terminal time can be cast in the minimisation of a cost function of the form Eq. (5). Throughout this study, the running cost has the form 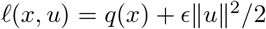, that is a trade-off between the squared magnitude of the control and a state-dependent cost measuring the squared distance from a target, *ξ, q*(*x*) = ||*x* – *ξ*||^2^/2.

For the class of cost functions in the form of Eq. (5), it is possible to solve the optimal control problem via dynamic programming. This is achieved by maximising, at every state *x*, the value function *J_u_*, defined as the negative of the cost-to-go function

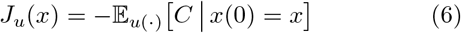

i.e. the cost to be paid conditioned on the initial state, averaged over all the realisations dynamics in Eq. (4), with control function *u*.

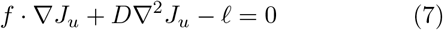

where *D* = *σ*^2^/2 and ∇ is the gradient with respect to the state variables *x*.

The value function corresponding to the optimal control *u*\*, denoted *J* * ≡ *J_u_*\*, therefore satisfies

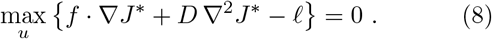

This equation, known as the *dynamic programming* (or *Bellman*) equation [41, 42], yields the optimal cost as well as the optimal control as a function *u*\* of the state *x*. The non-linearity introduced by the max operator, along with the infinite number of states (for continuous states and actions), makes the exact solution of Eq. (8) generally impossible.

Numerical techniques can be employed to find approximate solutions: reinforcement learning (RL) [24] with function approximation through deep neural networks [25, 43] is the numerical scheme used in this work for the solution of Eq. (8) for the optimal control of the ventral neural tube GRN. However, the case where *σ* is constant while *f* and *ℓ* have, respectively, linear and quadratic dependence on *u* (as in the case of the control in a landscape dealt with in the main text), falls into a general class of linearly solvable control problems [44, 45], in that Eq. (8) can be cast into a linear form through a change of variables (as detailed in SI, Sec. SI-1).

## SUPPLEMENTARY INFORMATION

### SI-1. Optimal control in a potential

Let us consider the Langevin dynamics

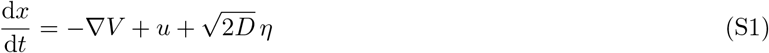

where *V* is a confining potential, *η* is a Gaussian noise with 〈*η*(*t*) *η*(*t*’)) = *δ*(*t* – *t*’) and *u* is an additional control drift. The control *u* is chosen to minimize a given cost functional, as detailed in the following. We choose the potential *V* in such a way that the uncontrolled dynamics has two stable fixed points (i.e. minima of *V*) at *x* = ±1: *V* (*x*) = *x*^4^/4 – *x*^2^/2.

#### a. Stationary-state optimization

We introduce the cost function

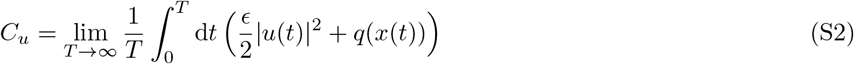

with

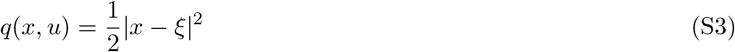

We seek to find the control strategy *u* that minimizes the expectation value of *C_u_* over all realisations of the stochastic dynamics Eq. (S1). If the system is ergodic, 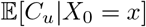 is a constant, i.e. it does not depend on the initial condition. In particular, this average is equivalent to that of the running cost at the stationary state:

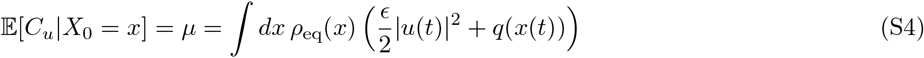

We can introduce the value function

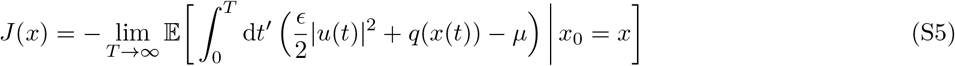

that is (minus) the *excess* cumulated cost from a given state relative to the steady state average. We can use the Feynman-Kac formula [46], to show that this satisfies

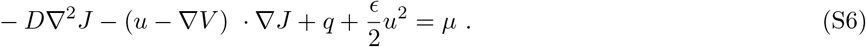

It can be verified by multiplying by the steady state (equilibrium) distribution *ρ*_eq_, satisfying (*u* – *VV*)*ρ*_eq_ = *DVρ*_eq_, and integrating over all states. The principle of dynamic programming holds that in order to minimize *μ*, it is sufficient to minimize *J*(*x*) for every *x*. We therefore see that the minimum condition for *J* yields

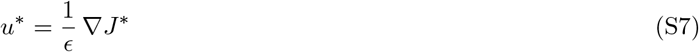

and that the optimal value function *J*\* satisfies the Bellman equation

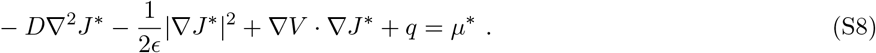

The constant *μ*\* is the minimum average cost at the stationary state. By replacing *J*\* = *ϵ*(*V* + 2*D* log *ψ*) this rewrites

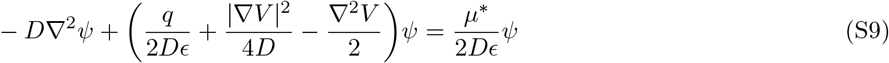

This is formally equivalent to the ground-state problem of a quantum particle of mass *m* = 2*D*/*ℏ*^2^ in the potential

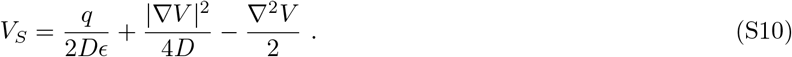

The change of variables implies that the optimally controlled dynamics is given by

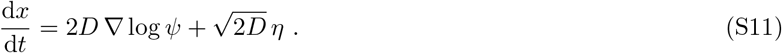

From the Fokker-Planck equation associated to Eq. (S11),

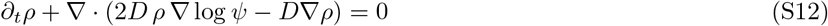

we see that the function *ψ* is related to the equilibrium steady-state distribution, *ρ*_eq_ ∝ *ψ*^2^. This ground-state problem can be solved by introducing a fictitious dynamics in imaginary time,

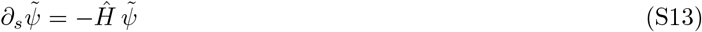

with the Hermitian operator *Ĥ* = –*DV*^2^ + *V_S_*. The ground state *ψ*_0_ of the Hamiltonian *Ĥ* is the slowest mode in the imaginary time evolution, and in the long-time limit, Eq. (S13) is solved by

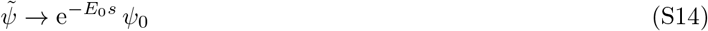

The solution of the HJB equation, *ψ*, then identifies with 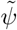, up to a scaling factor which depends solely on time. From the rate of change of the norm of 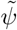 we can infer the minimum average cost:

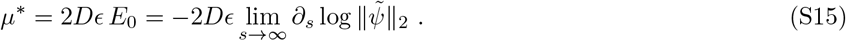

#### b. Exponential discounting

The control can also be chosen to minimize a cost over a shorter window of time, rather than at the steady-state. This can be done by introducing an exponential discount factor over time, as in

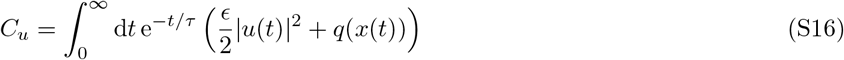

where *τ* sets a typical time scale over which rewards are accumulated in the future. As in the above case, we seek *u* that minimizes the expectation value 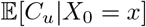 over the stochastic dynamics.

We can introduce the value function as (minus) the expected discounted cost-to-go from a given state at a given time

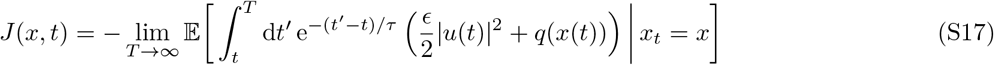

We see that this satisfies

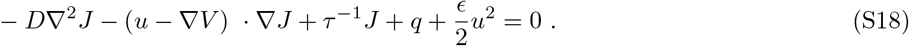

The optimality condition requires the control to be given by *u*\* = *ϵ*^1^∇*J*, and optimality Bellman equation writes

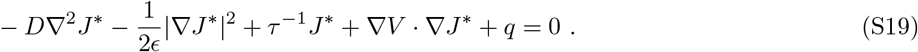

Analogously to the above case, with the transformation *J* * = *ϵ*(*V* + 2*D* log *ψ*), the Bellman equation takes the form

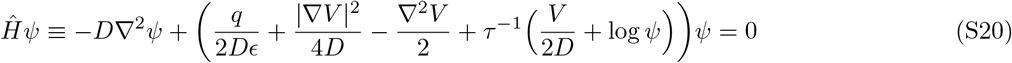

This non-linear Schrödinger equation can be solved numerically in a similar way as above, by introducing a fictitious dynamics in imaginary time, Eq. (S13), and solving it until convergence to the stationary state 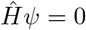.

#### c. Terminal cost and discounting

For a process that terminates with a probability per unit time *τ*^-1^ (or, in other terms, the probability density function for the terminal time is exponential, with mean *τ*), the exponential discount factor corresponds to the probability that a process that started at time t has not yet terminated at time *t*’:

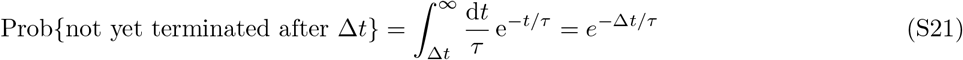

Therefore, the average of the cost *C_u_* in Eq.(S16) is equivalent to that of

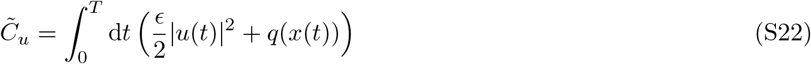

where *T* is the exponentially-distributed terminal time with mean *τ*.

For the dynamics with a terminal state (time), we can include a terminal cost at the time *T*, *Q*(*x*(*T*)). This is particularly relevant in the case of the cell-fate decision or the patterning example considered in the main text.

We can change the definition of the value function in Eq. (S17) by subtracting the contribution from the terminal cost. This can be written as

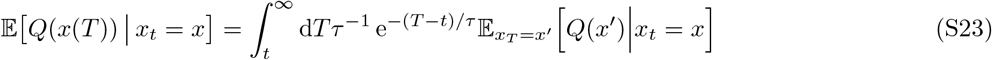

Together with the expression in Eq. (S17), the value for the task including the terminal cost can be expressed as

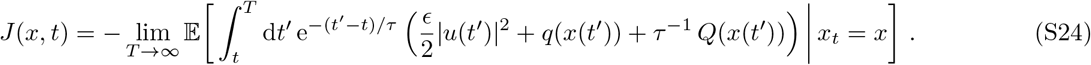

Therefore, we recognise that the addition of the terminal cost is equivalent to the replacement of the state-dependent running cost *q* by 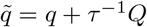 in Eq. (S16).

If we choose the terminal cost to be given by the same function *q* (the dimensions do not match, so we understand that *Q* is equal to q multiplied by a unit time constant), then 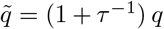. Since the optimal solution is invariant upon multiplications of the cost function by a global constant (see Eq. (S7)), the problem is equivalent to the one where *q* is kept the same, but *τ* enters as a rescaling of the trade-off parameter *ϵ*, replaced by 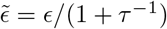.

#### d. First passage time near target

The mean first passage time (MFPT) at a given point 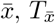 for a process starting at a point 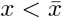, is expressed as

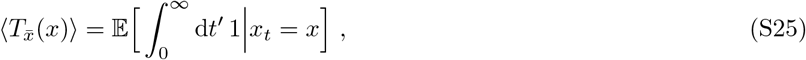

where the region 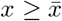 is replaced by absorbing states (viceversa if 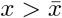). For the optimally control dynamics given in Eq. (S11), this satisfies [46]

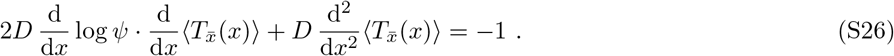

Its solution can be found by explicit quadratures, with the boundary conditions 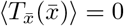 and 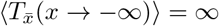,

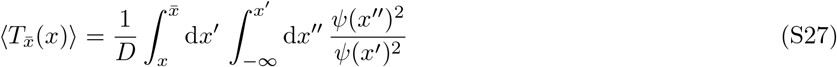

By interpreting *ψ*^2^ = exp(–*V*_eff_/*D*), we have

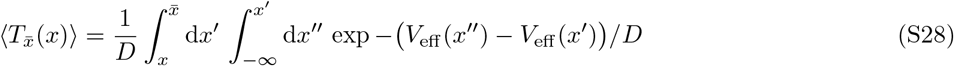

When *V*_eff_ has two minima, in the small-*D* limit, Eq. (S28) recovers the Freidlin-Wentzel theory of stochastic transitions via the saddle-point approximation [46, 47].

##### Low control and diffusion limit

For small values of *u*, the controlled potential *V*(*x, u*) still has two minima, corresponding to the stable fixed points of the controlled dynamics. If *D* is also small, the transitions between the two fixed points are rare, while typical realisations of the noise will produce small fluctuations around these: in this limit, Eq. (S28) gives the Freidlin-Wentzel theory of stochastic transitions [47], where the MFPT from the left minimum *x*_ to the right minimum *x*_+_ is therefore approximated as

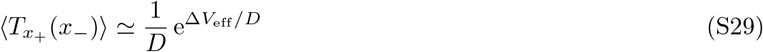

where Δ*V*_eff_ = *V*_eff_ (*x*_0_) – *V*_eff_ (*x*_), with *x*_0_ denoting the local maximum of the potential (or saddle) between the two minima. The rate for the opposite transition is analogously given by swapping *x*_ ↔ *x*_+_.

The steady-state probability to be near one or the other fixed point is given by the average exit time from the fixed point attractor. In the present example, this can be calculated as the MFPT from *x*_ ≃ – 1 to *x*_+_ ≃ 1, and vice versa.

First of all, we need to solve for the stationary points at a given value of *u*. In the linear approximation in *u*, these are

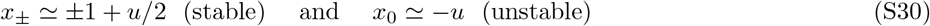

The value of the potential at these points is

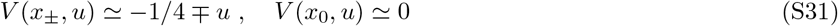

The MFPT for the “reverse” transition, 〈*T_x_*_ (*x*_+_)〉, and the MFPT for the “forward” one, 〈*T*_*x*+_ (*x*_)〉, are given by Eq. (S29), and their ratio gives the relative probability to be in the right or the left attractor at steady state:

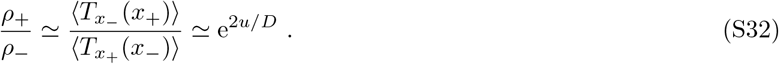

Therefore, we see that when *D* ≪ 1, for a range of control in the regime *D* ≪ |*u*| ≪ 1, the probability distribution is highly skewed towards one of the two attractors.

### SI-2. Environment dynamics

#### a. Ventral neural-progenitor GRN model (PONI network)

We outline here the details of the GRN model first presented in [13], with the addition of noise through the chemical Langevin equation approximation [8, 22].

We denote by *H*^+^ the Hill function

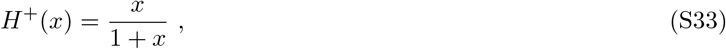

and by the latin letters the concentrations of the transcription factors, i.e. *P* ≡ [Pax6], *O* ≡ [Olig2], *N* ≡ [Nkx2.2], *I* ≡ [Irx3], *A* ≡ [GliA], *R* ≡ [GliR]. The dynamics of the four genes in the ventral neural tube GRN is described by the following system of first order ODEs:

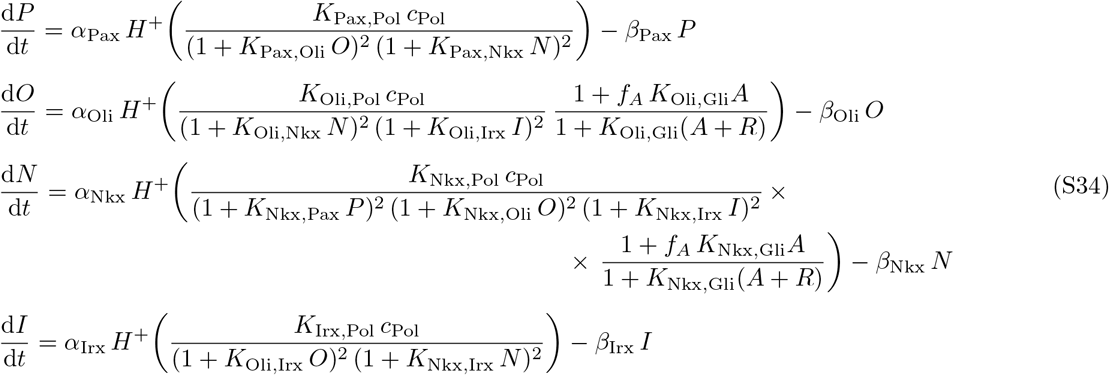

where *K_X,Y_* is the binding affinity of the TF/species Y onto its site on gene *X, f_A_* is the binding cooperativity factor for Gli activator, *c*_Pol_ is the (constant) concentration of RNAp, *α_X_* are the maximum production rates, and *β_X_* the degradation rates.

As in [8], we add (multiplicative) noise via the chemical Langevin equation (CLE) approximation [22] to the righthand side of Eqs. (S34). The overall size of the fluctuations is controlled by the inverse system size parameter, Ω^-1^. For instance, for Pax6, the multiplicative noise is modelled by

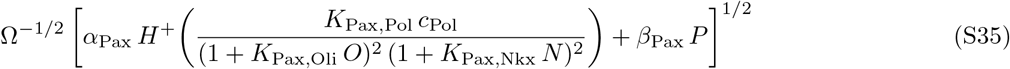

(i.e. the sum of production and degradation rates for the gene of interest, scaled by the inverse system size, under square root) multiplied by a standard Gaussian white noise, independent for each gene.

See Table I for the parameter values used.

**TABLE I.**
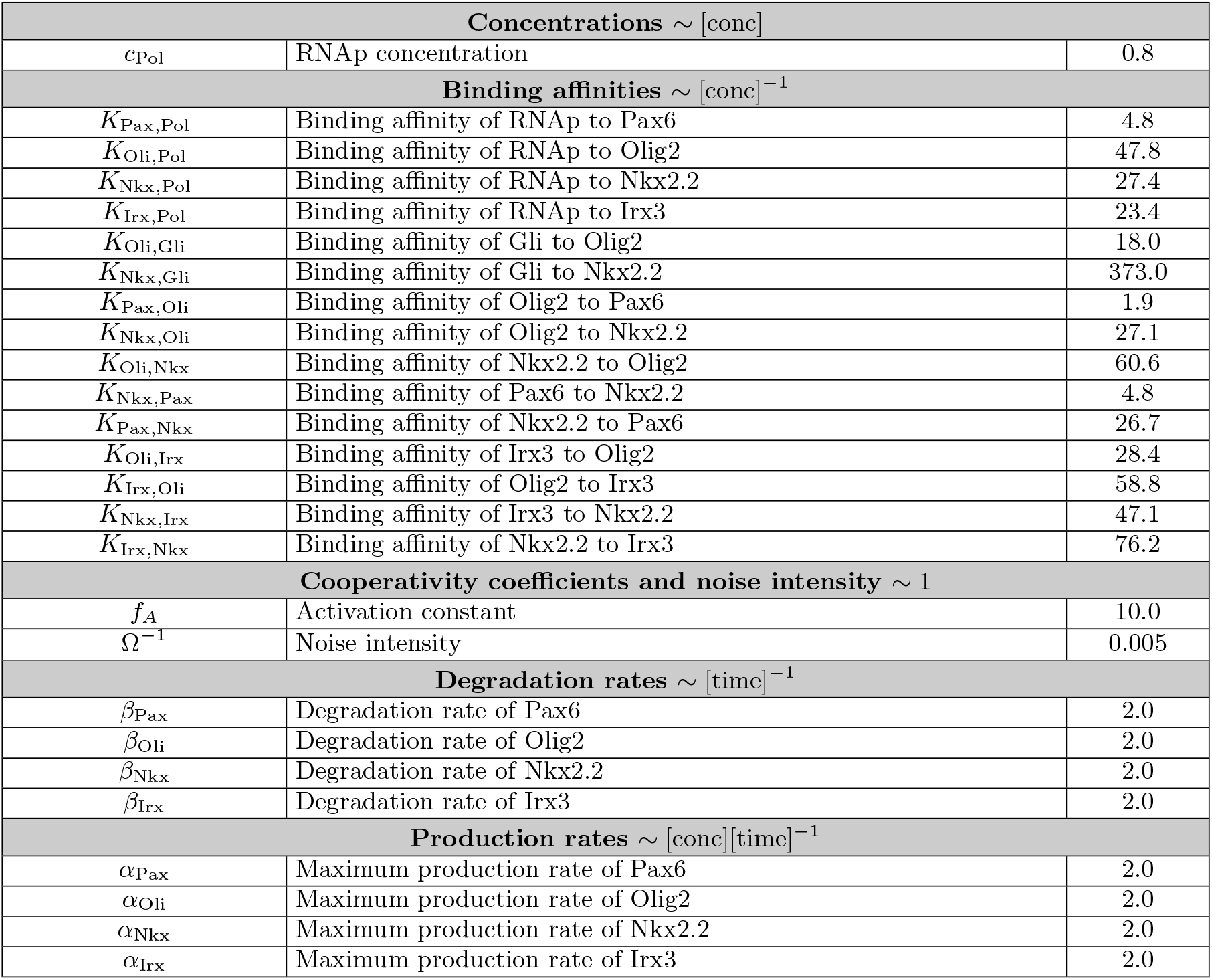
Parameters of the GRN model. Dimensionality of the constants are indicated in the header to every section.

| Concentrations $\sim [\text{conc}]$ | | |
| --- | --- | --- |
| $c_{\text{Pol}}$ | RNAp concentration | 0.8 |
| Binding affinities $\sim [\text{conc}]^{-1}$ | | |
| $K_{\text{Pax,Pol}}$ | Binding affinity of RNAp to Pax6 | 4.8 |
| $K_{\text{Oli,Pol}}$ | Binding affinity of RNAp to Olig2 | 47.8 |
| $K_{\text{Nkx,Pol}}$ | Binding affinity of RNAp to Nkx2.2 | 27.4 |
| $K_{\text{Irx,Pol}}$ | Binding affinity of RNAp to Irx3 | 23.4 |
| $K_{\text{Oli,Gli}}$ | Binding affinity of Gli to Olig2 | 18.0 |
| $K_{\text{Nkx,Gli}}$ | Binding affinity of Gli to Nkx2.2 | 373.0 |
| $K_{\text{Pax,Oli}}$ | Binding affinity of Olig2 to Pax6 | 1.9 |
| $K_{\text{Nkx,Oli}}$ | Binding affinity of Olig2 to Nkx2.2 | 27.1 |
| $K_{\text{Oli,Nkx}}$ | Binding affinity of Nkx2.2 to Olig2 | 60.6 |
| $K_{\text{Nkx,Pax}}$ | Binding affinity of Pax6 to Nkx2.2 | 4.8 |
| $K_{\text{Pax,Nkx}}$ | Binding affinity of Nkx2.2 to Pax6 | 26.7 |
| $K_{\text{Oli,Irx}}$ | Binding affinity of Irx3 to Olig2 | 28.4 |
| $K_{\text{Irx,Oli}}$ | Binding affinity of Olig2 to Irx3 | 58.8 |
| $K_{\text{Nkx,Irx}}$ | Binding affinity of Irx3 to Nkx2.2 | 47.1 |
| $K_{\text{Irx,Nkx}}$ | Binding affinity of Nkx2.2 to Irx3 | 76.2 |
| Cooperativity coefficients and noise intensity $\sim 1$ | | |
| $f_A$ | Activation constant | 10.0 |
| $\Omega^{-1}$ | Noise intensity | 0.005 |
| Degradation rates $\sim [\text{time}]^{-1}$ | | |
| $\beta_{\text{Pax}}$ | Degradation rate of Pax6 | 2.0 |
| $\beta_{\text{Oli}}$ | Degradation rate of Olig2 | 2.0 |
| $\beta_{\text{Nkx}}$ | Degradation rate of Nkx2.2 | 2.0 |
| $\beta_{\text{Irx}}$ | Degradation rate of Irx3 | 2.0 |
| Production rates $\sim [\text{conc}][\text{time}]^{-1}$ | | |
| $\alpha_{\text{Pax}}$ | Maximum production rate of Pax6 | 2.0 |
| $\alpha_{\text{Oli}}$ | Maximum production rate of Olig2 | 2.0 |
| $\alpha_{\text{Nkx}}$ | Maximum production rate of Nkx2.2 | 2.0 |
| $\alpha_{\text{Irx}}$ | Maximum production rate of Irx3 | 2.0 |

#### b. Dynamics of a stochastic gradient

In the patterning task, we also include a dynamics for the morphogen gradient. We simulate a non-stationary stochastic field 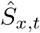, as the empirical number density field 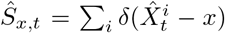 associated to a stochastic reactiondiffusion with

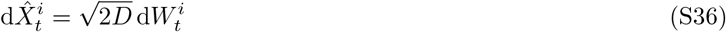

and where particles are removed with independent rates *κ* and added at *x*_0_ with rate *J*_0_. The SDE in Eq. (S36) provides an explicit method to simulate the spatio-temporal dynamics of the stochastic field 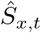. To do so, we simulate trajectories of Eq. (S36) via, e.g. Euler-Maruyama method, with time discretisation d*t*, that is

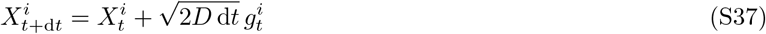

with 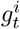 a normal-distributed random number with mean 0 and covariance 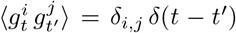; in the time step between *t* and *t* + d*t*, each particle is eliminated with probability *κ* d*t*, and a burst of *n_b_* new particles is added at *x*_0_ < 0 with probability *J*_0_ d*t*/*n_b_* (so that *J*_0_ is the overall average production rate, but with burst size *n_b_*). The number density field can be then defined with a spatial resolution d*x*, as the count of the number of particles within [*x* – d*x*/2, *x* + d*x*/2], divided by d*x*. The resolution d*x* is chosen to be the single-cell size.

We set the parameters of the dynamics as follows. 81 cells are aligned along one axis within [0,1], so d*x* = 1/80. The time discretization d*t* is chosen as 5 times smaller than that for the PONI network, but configurations are taken every 5 steps. The free parameters of the dynamics must set a time scale, a length scale and a typical number of particles. We set the overall time scale of the process through the degradation rate *κ*. The length scale is the decay length *λ* of the average gradient profile at steady state, 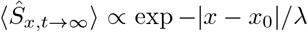. This is fixed to 0.15 in all simulations, consistently with experimental measures [16]. This decay length can be derived analytically to be 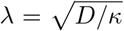, from which we fix the diffusion constant accordingly to be *D* = *κ λ*^2^. The typical density is chosen to be the average number density at *x* = 0 at steady state, which is 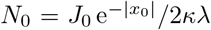. With a fixed burst rate *r* = *J*_0_/*n_b_* = 50, we modulate the burst size *n_b_* by inverting the expression for *N*_0_.

The ensemble average of the field 〈*S*〉, satisfies the PDE

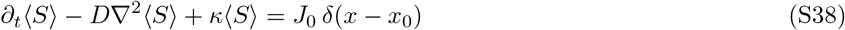

By integrating the spatial part, we can write

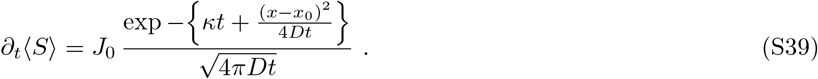

In Eq. (S39), the spatial variable enters only parametrically and the dynamics can be described as an ODE with time-dependent production rates. Therefore, (ensemble) averages of the signal experienced at different spatial locations can be regarded as “independent”, but at the expense of allowing non-autonomous dynamics for the local signal.

Parameters used for the simulations in this work are *λ* = 0.15 (in units of D-V axis length), *κ* = 0.5 (equal to *β*/4 – See Tab. I), and *N*_0_ = 5000.

### SI-3. Multi-Agent control

Here we derive the Bellman equation for the multi-agent (MA) case. The equations are written for the discrete-time and discrete-state case – as it is more transparent for a reinforcement learning implementation – but are easily generalized to continuous space and/or time. The notation is as follows:

- cell index, *i* (the 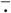 notation indicates arrays indexed by cells)
- cell state, including gene expression and extracellular signal levels, 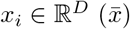
- target expression, 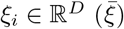
- intracellular signal, 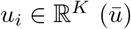
- M-A policy, 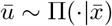, where 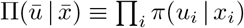
- model of the environment, 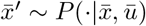

#### a. Full multi-agent case

The multi-agent probability distribution at time *t*, 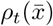, satisfies the forward Kolmogorov equation

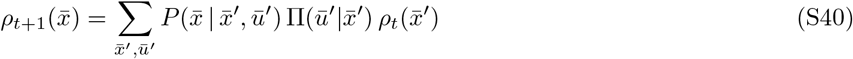

The goal of the agents is to maximize the expectation value of the discounted return (in the decision-making and reinforcement learning literature, it is more customary to express the goal in terms of maximisation of rewards, rather than minimisation costs):

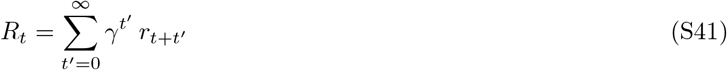

with

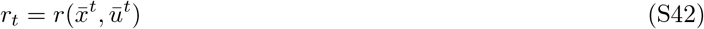

In the end, we will be interested in a reward of the form

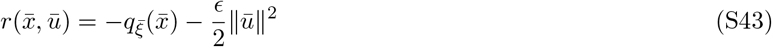

where, e.g. 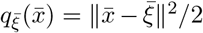. This negative reward is a cost that penalises certain configurations of the MA system -implementing the requirement to reach the target- and high values of control.

The objective function 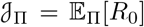, that is the ensemble average of *R*_0_ over the trajectories generated by the policy Π, writes

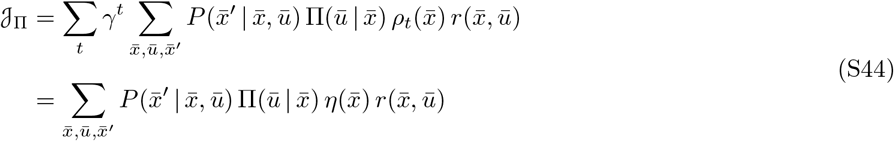

where *η* is the discounted occupancy

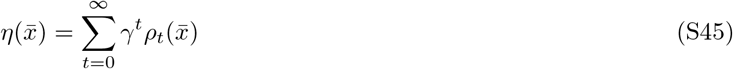

We can introduce the *quality* (or *state-action value)* function, which is the expectation value of the return conditioned on the initial state and action, 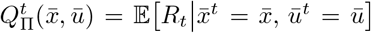. We can write a recursive equation of the value function 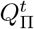, expressing the conditional expectation value 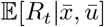 by making use of Eq. (S40):

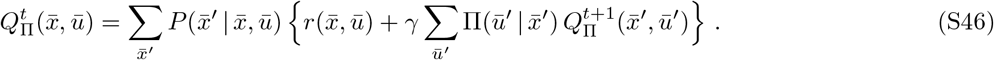

Since there is no finite horizon and neither rewards nor transition probabilities depend explicitly on time, we can seek for a stationary solution 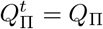.

The principle of dynamic programming [11, 48] consists in maximizing the expected return –i.e. the objective function 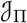– by maximizing its conditional expectation at intermediate times, that is the value function. The optimal policy Π*, then, is given in terms of the quality function as

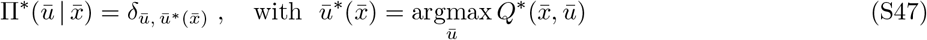

where the optimal quality function satisfies the Bellman equation

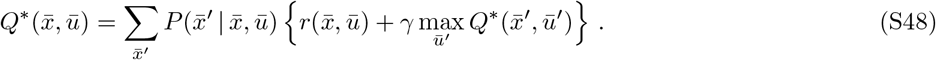

#### b. Independent agents

To reflect the requirement of each agent individually to reach their own target, we write 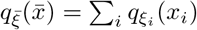, where *qξ* is some convex function that has a minimum at *ξ*. This is true for the cost rate 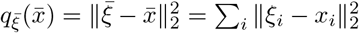.

So, the instantaneous reward for the MA system is the sum of rewards for the individual agents, *c_i_*, that are functions of the single agent’s observations and actions:

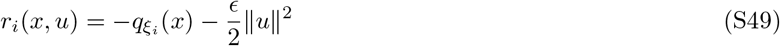

As discussed above, the MA policy Π with respect to which we want to optimize the performance is of the form

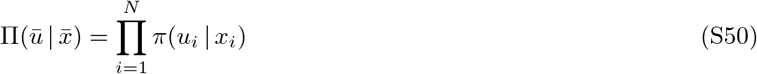

that is, actions by individual agents are chosen independently according to the same single-agent policy *π*. We seek for solutions of the Bellman equation of the form

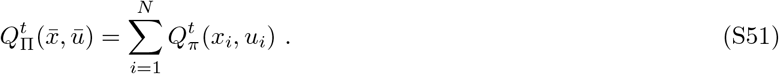

By replacing Eqs. (S50) and (S51), into the Bellman equation (S46), we have

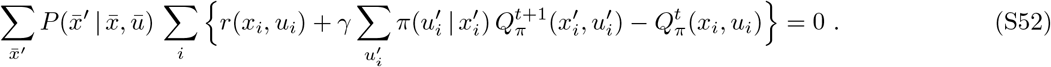

Optimality, in this approximation, is

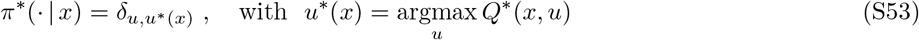

where *Q*\* denotes the optimal quality function solving

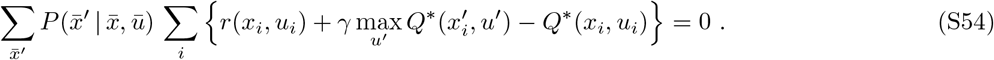

This is approximately solved by minimizing the expectation of the square MA error

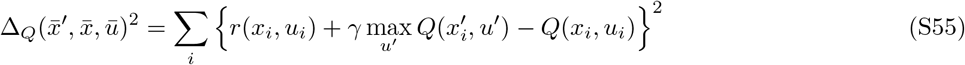

with respect to the *Q*,

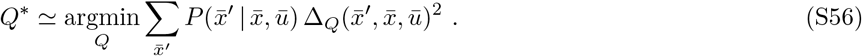

#### c. Memory in signal interpretation

The independent-agent ansatz is exact when the transition probabilities 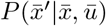 can be factorized into single-agent transition probabilities

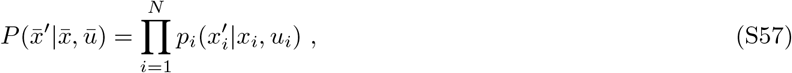

that is, when the dynamics of each agent is independent. This can be seen intuitively for a static and deterministic gradient. In such case, the (constant) value of the morphogen signal at the location of a given cell enters as a parameter in the quality function *Q*: it’s role is to “select” the specific single-agent problem for that particular cell. This effectively makes the MA task trivially decomposed into single-agent ones. If the gradient is stochastic and with a small noise, we could argue that the same holds in a probabilistic sense when the morphogen is at steady state or reaches it very fast (high *κ*). In general, when the morphogen gradient is modelled as a diffusion-degradation process –as in this case– this approximation is not valid. One can show that the average of the concentration field over the noise, 〈*S*〉, can be calculated as the solution of independent differential equations with local time-dependent rates (see Eq. (S39)). So, even though we may be able to express the average dynamics of the morphogen at individual cells locations as independent, 1) fluctuations will anyway be correlated and 2) we do so at the cost of introducing time dependence.

Here, we assume that it is possible to approximate the transition probability *P* by a factorized form as in Eq. (S57), at the expense of introducing auxiliary variables 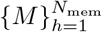, included in the “state” of the single cell along with its gene expression *G* and the local morphogen signal *S*. These memory variables integrate over time the extracellular signal *S* and that model the effective memory. We model these as the species in a signalling cascade, whereby *S* directly influences the production of *M*_1_, which in turn affects production of *M*_2_ etc.,

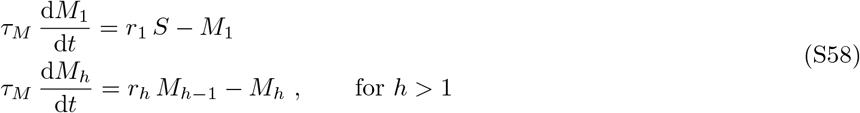

where *S* is the local morphogen concentration, and *r_h_* are components of the control vector *u*, therefore functions of the single cell state variables – bound between ±1. We choose the overall time constant *τ_M_* = 1. Notice that the dependence of the production rate for the memory variable *M_h_* depends at least linearly on *M*_*h*-1_: therefore, the control can modulate the production rates of the memory variables, but cannot be arbitrarily large for small signals.

### SI-4. RL solution

The approximate solution of Eq. (S52) via reinforcement learning (RL) requires the sampling of the tuples 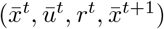. State-of-the-art deep-RL algorithms — such as DQN [43], DDPG [49], TD3 [25], SAC [50] etc— solve the problem of the stability of learning by storing a replay buffer 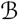 with the last *N*_replay_ tuples visited, and estimating gradients of the loss functions by averaging over a small number N_batch_ (batch size) of them.

Here we use TD3 [25], which is an actor-critic deep-RL algorithm, designed for continuous control problems. Similar to other actor-critic algorithm, it stores function approximators for both the policy (actor), and the value (critic) function. These are represented by deep neural networks with parameters *ϕ* and *θ*, respectively (*π* ≃ *π_ϕ_* and *Q* ≃ *Q_θ_*). In order to reduce the bias in the estimate of the value function *Q*, TD3 uses two critics *(T* for “twin”).^1^ As in other deep-RL AC algorithms, in order to make learning more stable, TD3 stores two copies of each function approximator: the first is updated on-line; the second is used as target and integrates the first at a slow rate, and with delay. TD3 uses a SARSA-like target for the value function, by sampling the next action using the target policy.

We here use the TD3 algorithm for episodic tasks (see [25] for details). We use *α* = 10^−3^, *β* =10^−3^. All other details are the same as in the original paper. The discount factor (which is a property of the task!) *γ* = 0.99, which for time step d*t* = 0.005 corresponds to the exponential discount time in continuous time *τ* ~ 5.

In the case of the MA problem described above, we need to modify this algorithm by storing transitions of the MA system, defining a target for each individual agent (based on their single-agent rewards, states and actions), and averaging gradients over the agents as well. This is detailed in Alg. 2. The learning rates here are *α* = 3 × 10^5^ and *β* = 10^-5^.

---

**FIG. S1.**
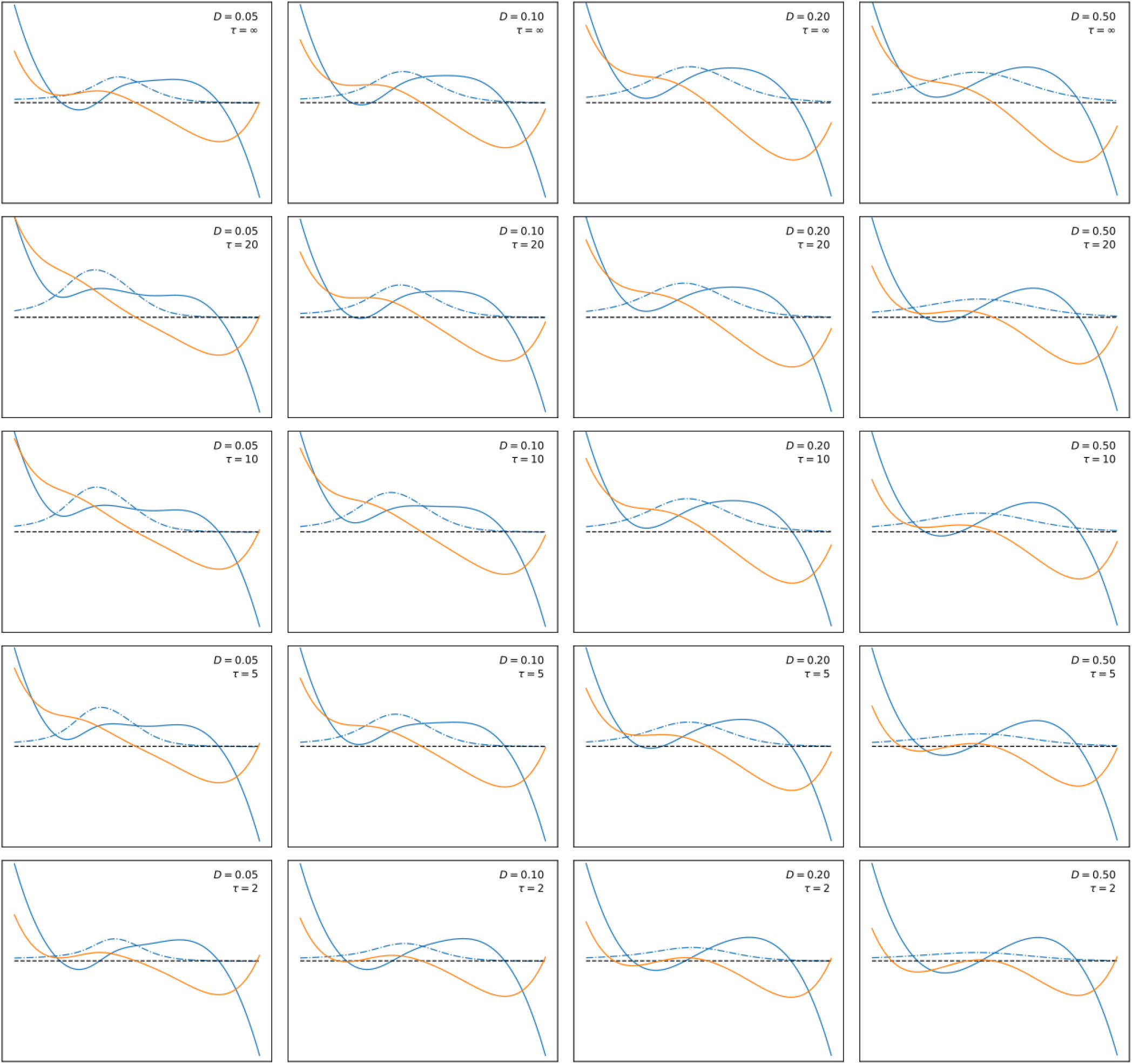
Optimally controlled flow (solid blue), optimal control (dashed-dotted blue) and landscape (solid orange), for an array of values of *D* and *τ*. The cost for control is set to *ϵ* = 10 in all panels.

**FIG. S2.**
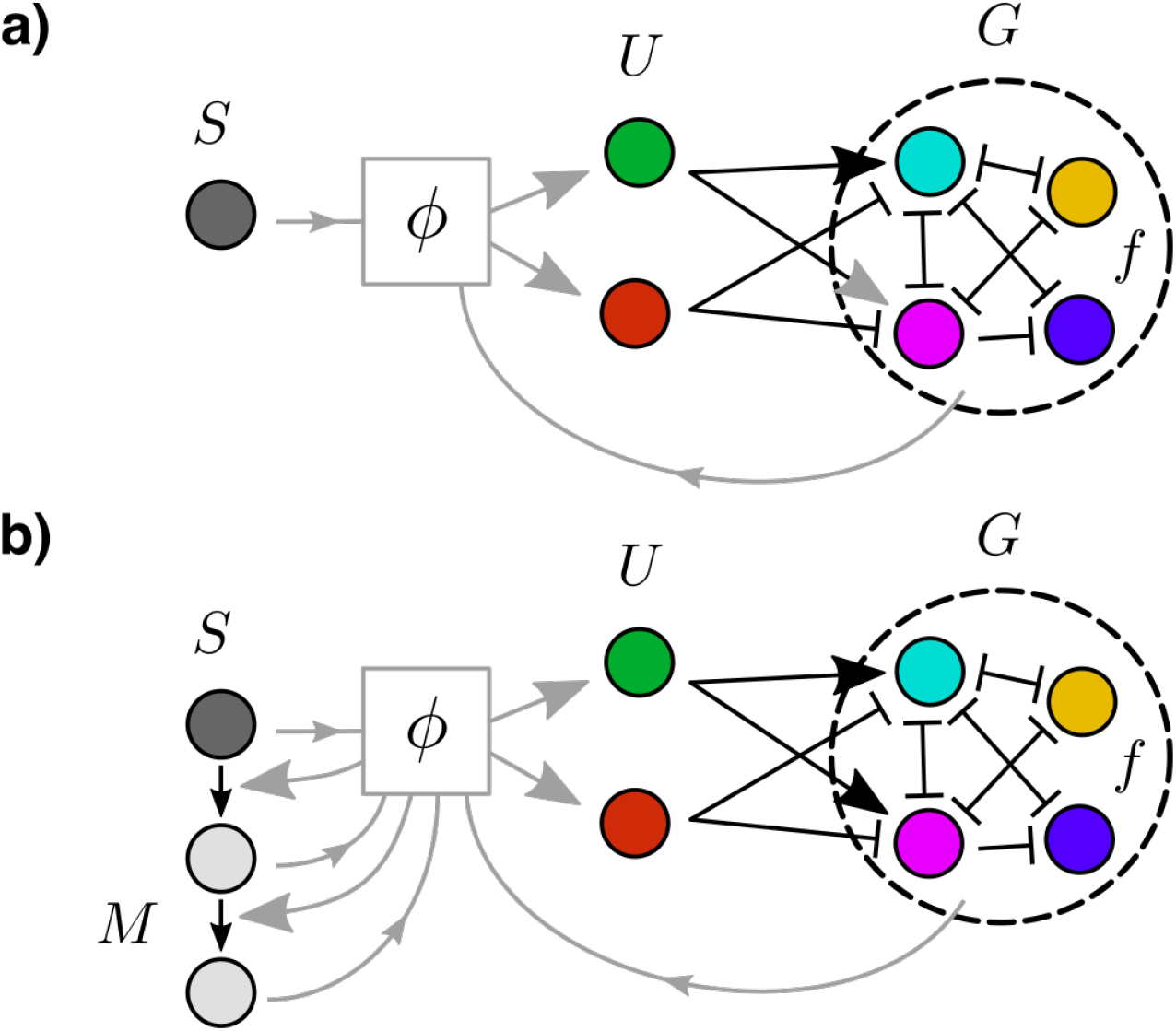
Scheme of the model of the environment. The model where the local morphogen signal is added to the GRN concentration to give the full state of the environment (a) is augmented by adding variables –in this case 2– that integrate the signal and contain memory information (b).

**FIG. S3.**
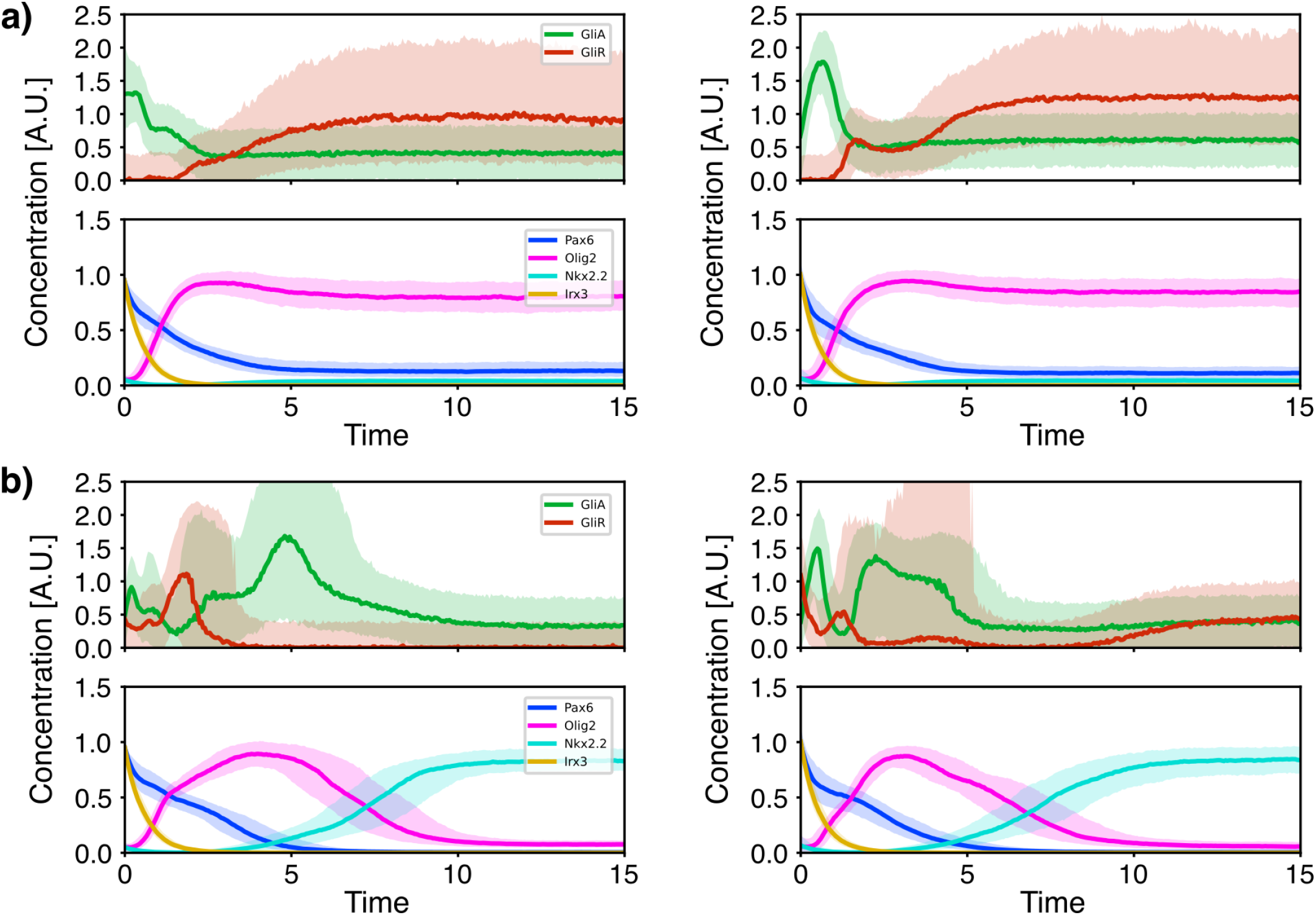
Comparison between different reinforcement learning solutions for the optimal control of the ventral neural tube GRN [13]. The solution presented in the main text (left) compared with the best solution of a different experiment with the same algorithm (right), for (a) the Olig2+ target and (b) the Nkx2.2+ target.

**FIG. S4.**
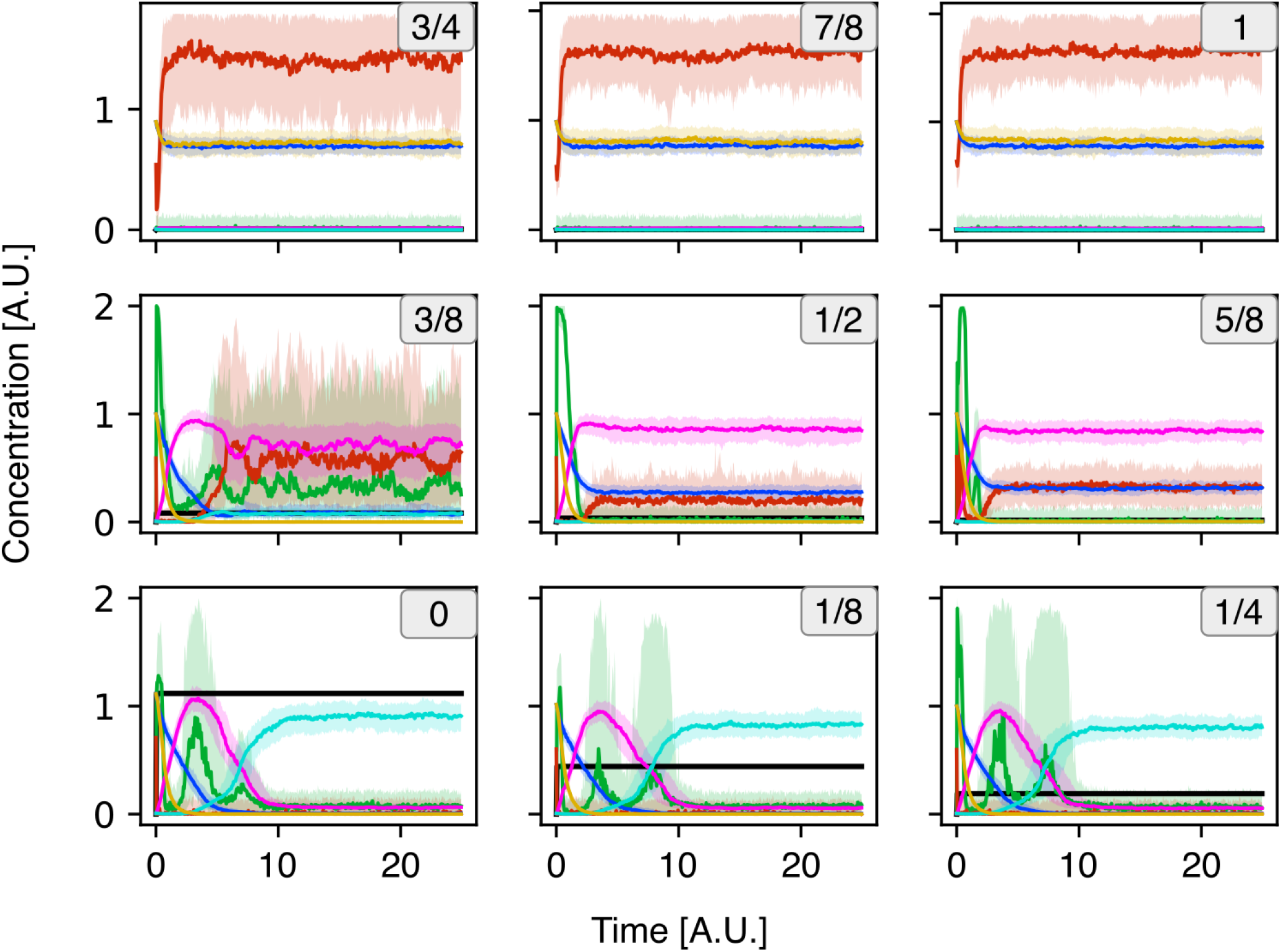
Patterning dynamics for static gradient, when the independent agent ansatz is exact

## Footnotes

1 In standard Q-learning, the value of the state after the transition is taken to be the maximum over all actions of the *Q* function evaluated at that state, by boostrapping. This is a problem that is present also in actor-critic algorithms like DDPG, where the “maximization over actions” is implicit in the policy-gradient formula. This typically leads to an overestimation of the value (as demonstrated in the paper).

## Notes

### Competing Interest Statement

The authors have declared no competing interest.

